# Cell type diversity in a developing octopus brain

**DOI:** 10.1101/2022.01.24.477459

**Authors:** Ruth Styfhals, Grygoriy Zolotarov, Gert Hulselmans, Katina I. Spanier, Suresh Poovathingal, Ali M. Elagoz, Astrid Deryckere, Nikolaus Rajewsky, Giovanna Ponte, Graziano Fiorito, Stein Aerts, Eve Seuntjens

## Abstract

Octopuses are mollusks that have evolved intricate neural systems comparable with vertebrates in terms of cell number, complexity and size. The cell types within the octopus brain that control their amazingly rich behavioral repertoire are still unknown. Here we profile cell diversity of the paralarval *Octopus vulgaris* brain to build a comprehensive cell type atlas that comprises mostly neural cells, as well as multiple glial subtypes, endothelial cells and fibroblasts. Moreover, we spatially map cell types within the octopus brain, including vertical and optic lobe cell types. Investigation of cell type conservation reveals a shared gene signature between glial cells of mice, fly and octopus. Genes related to learning and memory are enriched in vertical lobe cells, which show molecular similarities with Kenyon cells in *Drosophila*. Taken together, our data sheds light on cell type diversity and evolution of the complex octopus brain.

**Highlights & Key findings:**

- Characterization of different cell types present in the early paralarval brain
- Cross-species comparisons reveal a conserved glial gene expression signature
- Vertical lobe amacrine cells in octopus have molecular similarities to fly Kenyon cells
- Homeobox genes are defining transcription factors for cell type identity
- Recently expanded gene families may underlie cellular diversification

## Introduction

Cephalopods, such as cuttlefish, squid and octopus, are enigmatic organisms that have evolved impressive cognitive capabilities. They can display a range of complex behaviors like problem-solving, tool use and millisecond camouflaging skills, for which higher cognitive functions are likely required^1–4^. Although the basic design of an octopus brain seems typically invertebrate-like, with neuropil surrounded by a layer of monopolar neuronal cell bodies, its anatomical complexity is unparalleled among invertebrates. Octopuses have a large centralized brain with more than 30 differentiated lobes and an intricate organization to support the transfer, integration and computation of information^5,6^.

The octopus brain consists of: (1) two optic lobes that are involved in visual sensory processing and memory storage of visual information, (2) the supraesophageal mass, a sensory-motor, associative and integrative center, which contributes to long-term memory storage and (3) the subesophageal mass, responsible for motor and visceral coordination and other sensory processing^6^. Even if it is generally accepted that complex brains and intelligence arose multiple times during evolution^3,7,8^, the necessary building principles to create complex brains remain unknown. Perhaps the most intriguing part of the octopus brain is the vertical lobe, with its 26 million neurons^5^. The vertical lobe has been posited to be the functional analog of the invertebrate mushroom body and the mammalian pallium^9,10^.

The central nervous system of an adult octopus consists of 200 million cells, which is comparable with the number of neurons in the brain of a tree shrew^5,11^. Although cell types present in the adult octopus brain have been extensively characterized morphologically by J.Z. Young^6^, the molecular signature of these cell types, and whether such a signature is similar to certain cells in the fly or mouse nervous system, remains unresolved.

Since the first octopus genome was published in 2015^12^, there has been increasing interest in cephalopod biology and neuroscience. This pivotal study highlights the expansion of gene families such as protocadherins (PCDH), G-protein coupled receptors (GPCR) and Zinc-finger transcription factors (ZnF) in *Octopus bimaculoides*^12^. These gene families have known roles in brain development and neural wiring in complex-brain species^13–15^. Whether these peculiar expansions have contributed to cell type diversity in octopus is unknown.

The common octopus, *Octopus vulgaris*, lays over ten thousands of transparent eggs, which take approximately one month to complete embryonic development and hatching^16^. At this point, the free swimming paralarvae undergo a planktonic phase before they adapt to a benthic lifestyle^17^. Their brain develops from the lateral lips - an embryonic neurogenic region surrounding the eyes - and contains all major lobes of the adult structure in miniature form^18,19^, see Fig. 1a-c. Upon hatching, the paralarval brain consists only of an estimated 200,000 cells^20^ and it is about four times the size of the adult fruit fly brain, which makes it an attractive structure to build a cell type atlas. Since the development of single-cell RNA sequencing technologies, most studies have been model organism oriented. Indeed, studying cell type diversity in the absence of known cell type gene expression signatures poses additional challenges.

**Fig. 1.**
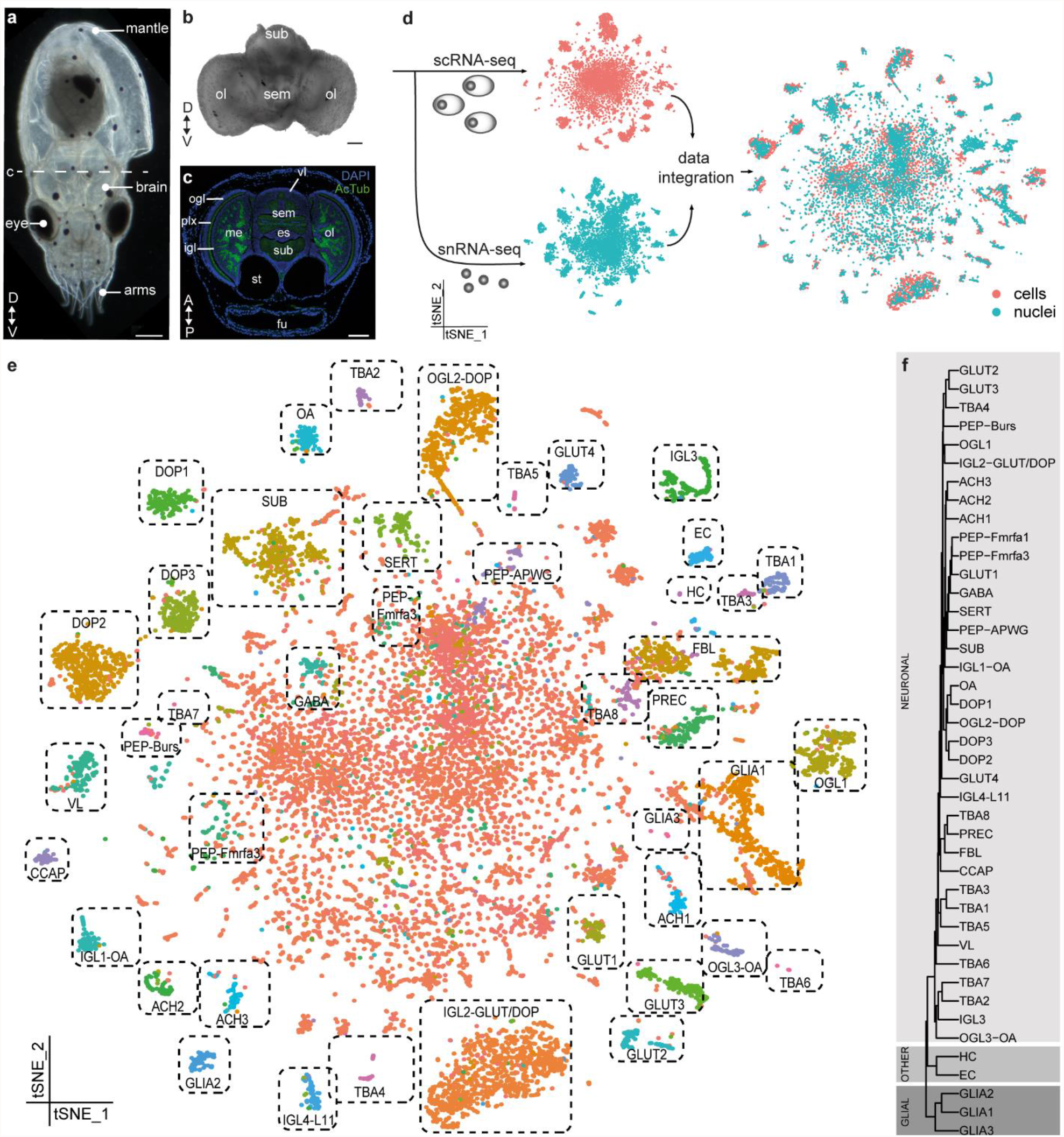
Cellular diversity in the developing octopus brain. **a** One-day-old *Octopus vulgaris* paralarva. Dashed line indicates the sectioning plane in c. **b** Dissected brain. **c** Representative transversal section of a larva with annotated anatomical structures. **d** Experimental design of this study. Single cell and nuclei RNA sequencing was performed with 10x Genomics and data integration resulted in a filtered dataset of 17,081 high quality cells. **e** t-SNE representation of the integrated sc and snRNA-seq data. Annotated cell types are labeled. **f** Hierarchical clustering of all stable clusters. All scale bars represent 100 µm. A, anterior; ACH, cholinergic neurons; AcTub, acetylated tubulin; CCAP, cardioactive peptide cells; DOP, dopaminergic neurons; D, dorsal; EC, endothelial cells; es, esophagus; FBL, fibroblasts; fu, funnel; GABA; GABAergic neurons; GLUT, glutamatergic neurons; HC, hemocytes; igl, inner granular layer; IGL, inner granular layer cells; me, medulla; OA, octopaminergic neurons; ogl, outer granular layer; OGL, outer granular layer cells; ol, optic lobe; P, posterior; PEP, peptidergic neurons; plx, plexiform layer; sem, supraesophageal mass; SERT, serotonergic neurons; sub, subesophageal mass; SUB, subesophageal neurons; st, statocysts; TBA, to be annotated; V, ventral; vl, vertical lobe; VL, vertical lobe cells.

In this study, we report for the first time on cell type diversity in the *Octopus vulgaris* brain. Comparing and combining single-cell and single-nuclei datasets, we systematically characterize 42 cell types within the larval brain of *O. vulgaris* and provide the first description of their transcriptomes. We spatially map several of these cell types with *in situ* hybridization and use cross-species comparisons to predict conserved cell types and compare gene expression signatures. We provide evidence that several cell types display unique combinations of PCDH, ZnF or GPCR, suggesting that octopus-specific gene expansions contributed to increased cell type diversity. While we estimate the diversity of octopus brain cell types to be larger than our current view, our results are a valuable resource for future physiological studies and offer novel insights into the molecular profile of octopus brain cells and the evolution of cell types.

## Methods

### Genome annotation

The chromosomal scale genome assembly for *Octopus sinensis* was used (ASM634580v1)^21^. We extended the 3’-ends of the genes using an evidence-guided approach. First, full isoform-sequencing data (Iso-Seq, PacBio Sequel) was used to reconstruct mRNA isoforms (data retrieved from PRJNA718058, PRJNA791920). We included both paralarval^18^ and adult^22^ Iso-Seq datasets of *O. vulgaris*. For each gene, the end of the longest isoform was considered as the new 3’-end. Next, a full-length mRNA sequencing method - FLAM-seq - was used to locate mRNA cleavage sites in the genome^22^. Cleavage sites located within 60,000 bp were assigned to the closest upstream genes (PRJNA791920). Finally, to account for the genes missing in the FLAM-seq dataset, published short-read RNA-seq datasets^23^ were used to extend the genes based on coverage. In brief, each gene was extended if there was sufficient continuous RNA-seq coverage (≥ 5 reads) downstream. A schematic depiction of the pipeline is available in Fig. S1C. The resulting genome annotation is available in Data S1. This approach resulted in a twofold decrease in the number of the reads mapping to intergenic regions (Data S2). We manually curated the genome annotation for the PCDH gene family. Some read-through transcripts resulted in gene fusions and this was corrected by taking into account the number of protein domains. Transdecoder (https://github.com/TransDecoder/, v5.5.0) was used to identify the CDS. Functional annotation was performed by running BLAST+ v2.7.1 against the SwissProt protein databases of *Drosophila melanogaster, Mus musculus* and *O. bimaculoides* (with a e-value threshold of 10^−5^). In addition, EggNOG-mapper v2^24^ was used to infer orthologies to bilaterian genes. The results are summarized in Data S3. Gene ontology terms were also predicted by EggNOG and we calculated the enriched gene ontology terms for certain clusters with the GSEApy package (v0.10.3).

### Animals

*O. vulgaris* embryos were obtained from the Instituto Español de Oceanografía (IEO, Tenerife, Spain). Embryos were then incubated until hatching in a closed system in the Laboratory of Developmental Neurobiology (KU Leuven), Belgium^16^. One day after hatching, larvae were sedated with 2% ethanol filtered artificial seawater. Next, 30 brains were dissected on ice for single cells and 30 brains for single nuclei in L15-medium (Sigma) with additional salts (214 mM NaCl, 26 mM MgSO_4_x7H_2_O, 4.6 mM KCl, 2.3 mM NaHCO_3_, 28 mM MgCl_2_x6H_2_O, 0.2 mM L-glutamine, 38 mM D-glucose, 10 mM CaCl_2_x2H_2_O, pH=7.6). Statocysts and retinal tissues were removed as much as possible. All procedures involving hatchlings were covered by animal ethics permit P080/2021 and in accordance with the European guidelines for cephalopod research^25^.

### Immunohistochemistry and *in situ* hybridization

One-day-old paralarvae were sedated in artificial seawater with 2% ethanol and fixed overnight in 4% paraformaldehyde (PFA) in phosphate buffered saline (PBS). Immunohistochemistry and *In situ* hybridization were performed as described before^18^. Briefly, embryos were embedded in paraffin after progressive dehydration and sectioned using a paraffin microtome (Thermo Scientific, Microm HM360) to obtain 6 µm-thick transversal sections. For immunohistochemistry, we used mouse anti-Acetylated alpha Tubulin (Sigma T6793) and rabbit anti-phospho-histone H3 (Ser10) (Millipore 06-570) as primary antibodies. Colorimetric *in situ* hybridization was performed using DIG-labeled antisense probes with the use of an automated platform (Ventana Discovery, Roche) with RiboMap fixation and BlueMap detection kits (Roche). The amount of probe used (100-300 ng) and incubation with BCIP/NBT (6-9 hours) was dependent on the target gene. Each probe was tested at least twice (different embryos and independent experiments). Probe sequences are listed in Data S6. Hybridization chain reaction (HCRv3.0) and imaging was performed as described before^18^. Probe sets were ordered for *Ov-glut, Ov-th, Ov-vacht, Ov-LOC118767670 and Ov-apolpp* from Integrated DNA Technologies, Inc (Data S6). Probe sets were designed with the insitu_probe_generator^26^ followed by automated blasting and formatting to minimize off-target hybridization with a custom script^27^. Imaging was done either with a Leica DM6 upright microscope (IHC, colorimetric ISH) or an Olympus confocal microscope Fluoview FV1000 (HCR).

### Single cell suspension

Paralarval brains were enzymatically dissociated by adding 20 µl of Collagenase/Dispase (100 mg/ml, Roche) to 500 µl L15-adapted medium and incubating for two hours at 25°C, 500 rpm. Every 15 minutes, a P100 was used to pipet slowly up and down until the tissue was fully dissociated. After a 5 min centrifugation step (200 rcf, 4°C), the supernatant was discarded and the pellet was resuspended in 1 ml of Mg-Ca-Free filtered sea water with 0.04% BSA (449 mM NaCl, 33 mM Na_2_SO_4_, 9 mM KCL, 2.15 mM NaHCO_3_, 10 mM Tris-Cl pH 8.2, 2.5 mM EGTA, filter sterilized). The cells were pulled through a strainer (35 µm) by a brief spin, followed by a wash with 400 µl Ca-Mg-Free filtered seawater. Cells were centrifuged again for 5 min (200 rcf, 4°C), supernatant was removed and the pellet was resuspended in 100 µl Ca-Mg-Free filtered sea water with 0.04% BSA. The cell viability and concentration were assessed by the LUNA-FL Dual Fluorescence Cell Counter (Logos Biosystems). We obtained a single cell suspension with a multiplet cell percentage of 2.6%. Average cell size was 9.1 µm. The cell suspension was further diluted to reach appropriate cell counts, and a final viability of 84.9% was obtained before proceeding with 10X Genomics.

### Single nuclei extraction

The brains were immediately transferred to a dounce homogenizer (Sigma) containing 0.5 ml of ice-cold homogenization buffer (HB) (320 mM Sucrose, 5 mM CaCl_2_, 3 mM Mg(OAc)_2_, 10 mM Tris 7.8, 0.1 mM EDTA, 0.1% IGEPAL CA-360, 0.1 mM Phenylmethylsulfonyl fluoride, 1 mM β-mercaptoethanol with 5 µl RNasin Plus). Tissue was incubated in the HB for 5 min before starting homogenization. The tissue was homogenized with 10 manual gentle strokes (pestle A) + 10 manual gentle strokes (pestle B). The tissue homogenate was filtered through a 70 µm cell mesh strainer. Leftover contents on the strainer were washed with an additional 0.5 ml HB buffer. The homogenized tissue was incubated in HB on ice for 5 min. Leftover contents on the strainer were washed with an additional 1.65 ml HB, which added to a final volume of 2.65 ml. The nuclei homogenate in the HB was mixed with 2.65 ml of Gradient Medium (GM) (5 mM CaCl_2_, 50% Optiprep, 3 mM Mg(OAc)_2_, 10 mM Tris 7.8, 0.1 mM Phenylmethylsulfonyl fluoride, 1 mM β-mercaptoethanol). 29% density cushion was prepared by dilution of Optiprep with Optiprep Diluent Medium (150 mM KCl, 30 mM MgCl_2_, 60 mM Tris pH 8.0, 250 mM sucrose). The nuclei suspension in the HB + GM mix was layered over the 29% cushion and centrifuged in an SW41Ti rotor (Beckman Coulter) at 7700 rpm and 4°C for 30 min. The supernatant was removed with a Pasteur pipette, and the removal of the lower supernatant was done with a P200. The nuclei pellet was resuspended in 50 µl Resuspension Buffer (PBS, 1% BSA) and transferred to a new tube. The resuspended nuclei were counted using a LUNA-FL Dual Fluorescence Cell Counter (Logos Biosystems).

### 10X Genomics

Library preparations for the sc/snRNA-seq experiments were performed using 10X Genomics Chromium Single Cell 3’ Kit, v3 chemistry (10X Genomics, Pleasanton, CA, USA). We aimed for a targeted cell recovery of 6000-10,000 cells/nuclei. Post cell count and QC, the samples were immediately loaded onto the Chromium Controller. Single cell or single nuclei RNA-seq libraries were prepared using manufacturers recommendations (Single cell 3’ reagent kits v3.1 user guide; CG000204 Rev D), and at the different check points the library quality was assessed using Qubit (ThermoFisher) and Bioanalyzer (Agilent). With a targeted sequencing coverage of 25-50 K reads per cell, single cell libraries were sequenced on Illumina’s NovaSeq 6000 platform (VIB nucleomics core, KU Leuven) using paired-end sequencing workflow and with recommended 10X; v3.1 read parameters (28-8-0-91 cycles). A total of 202,402,758 reads were obtained for the nuclei and 247,457,191 reads for the cells.

### 10x Data preprocessing

All samples were processed with 10x Genomics Cell Ranger 5.0.1 for mapping, barcode assignment and counting. Introns were retained and the parameter --expected cells was set at 8000 for both samples. Sequencing metrics for both samples can be found in Data S2. The 3’-end extended genome annotation described above was used as a reference (Data S1). This resulted in a raw dataset of 20,957 genes by 14,265 cells for the single cells and 21,073 genes by 8910 cells for the single nuclei. Filtering and subsetting steps were done in Seurat v3.2.3^28^. Nuclei and cells with too high (>4000) or too low (<400 for nuclei, <800 for cells) gene counts were filtered out. Cells with a higher percentage (>5) of mitochondrial RNA were regressed out. Genes expressed in less than 10 cells were excluded. The SCTransform scaling method was used and data integration of the cells and nuclei was done following the recommended Seurat vignette. This resulted in a filtered integrated dataset of 17,961 genes by 17,081 cells.

### Cluster annotation

The package scclusteval was used to assess optimal clustering parameters to obtain the highest number of stable clusters^29^. By resampling and repeated clustering, we used the mean Jaccard indices as a metric for stability. Reclustering according to these optimal parameters (dims =150, k.param =10, resolution =2) resulted in the highest number of stable clusters. Cluster identities were transferred to the Seurat object. Differentially expressed genes were calculated for all clusters compared to all other clusters (logfc.threshold = 0.25). We used the package SCopeLoomR (https://github.com/aertslab/SCopeLoomR; v0.13.0) to generate the loom file, to facilitate data exploration in SCope. The expression levels of the genes in the t-SNE plots are Log transformed and visualized with a scale bar. For cluster annotation purposes we filtered out all unstable clusters (<0.6 Jaccard index) and discarded the clusters that were not well defined (clusters 58,0,12,17,48,8). Cluster 3 and cluster 15 were merged into IGL2-GLUT/DOP. These two clusters largely overlapped and did not have many differentially expressed genes. We attributed this to a batch effect of the nuclei and cells. This resulted in a dataset of 42 robust clusters. Cell type annotation was based on the expression of vertebrate and invertebrate marker genes. Cell types were named based on their spatial localization and/or their neurotransmitter/neuropeptide phenotypes (Data S4). We used the PrctCellExpringGene function to calculate the % of cells that expresses a certain gene (number of cells with raw counts > 0).

### Transcription factors and cell type specificity

Transcription factors (TF) were annotated with animalTFDB^30^. Gene expression was averaged per cell type based on the SCT assay and genes expressed in less than 20 cells were excluded. The ComplexHeatmap R package was used for data visualization (Fig. 6). We calculated the tau value for all transcription factors using the tspex Python package (v0.6.2). We then calculated whether a transcription family was enriched within the rank of tau (GSEApy, v0.10.3).

### Cross-species cell type comparison

SAMap v0.1.6^31^ was used to compare our data to scRNA-seq datasets of different species to gain more information about the identity and evolution of the octopus cell types. We mapped the octopus paralarval brain to a mouse brain dataset ^32^ and to the adult fly brain^33^. Only alignment scores above 0.25 were considered to be of significance. Resulting annotations were visualized on the octopus t-SNE plot (Fig. 4) and listed in Data S4.

### Gene family enrichment analysis

Fisher’s exact test was performed to calculate statistical enrichment for these recently expanded gene families such as PCDH, C2H2-ZnF and GPCR. Contingency tables were constructed and we then compared the number of genes belonging to a certain gene family to all other genes present in that cell type versus all other cell types. Only genes with an avg_logFC of above 0.25 or below -0.25 were considered for this analysis (75 PCDHs, 141 C2H2-ZnF and 130 GPCRs). Fisher’s exact tests for each cell type were followed by a bonferroni correction for multiple testing with p.adjust() in R studio. Expression of octopus-specific genes was averaged per cell type based on the SCT assay and visualized on a scaled heatmap (Fig. S9). The ComplexHeatmap R package was used for data visualization and significant enrichments were highlighted in red (Fig. S9).

## Results

### Generation of a single-cell and single-nucleus transcriptome atlas

To comprehensively study cellular diversity in the octopus brain, we performed 10x Genomics on single cells and single nuclei from dissected brains of one-day-old *O. vulgaris* paralarvae (Fig. 1a). The octopus hatchling brain possesses about 200,000 cells and consists of two optic lobes (ol), the supraesophageal mass (sem) and subesophageal mass (sub) that surround the esophagus (Fig. 1b-c). We dissociated the brains to create a single cell suspension for single-cell RNA sequencing and extracted nuclei for single-nuclei RNA sequencing (Fig. 1d, see also Methods).

As the draft genome of *O. vulgaris* was too fragmentary for annotation^34^, we mapped the reads to the chromosomal scale genome assembly for *O. sinensis*, a very closely related species to *O. vulgaris*^21^. Furthermore, to optimize the accuracy of gene expression counts, we created an improved gene annotation. Since the single-cell RNA-seq method used here is biased towards 3’ ends of messenger RNAs, we focused on the 3’ UTR annotation (Fig. S1). Particularly, we used FLAM-seq^35^ and Iso-Seq (PacBio) full-length mRNA sequencing data of embryonic, paralarval and adult octopus tissue (Fig. S1, Data S1)^18,22^. With this new annotation, the percentage of reads that mapped confidently to the transcriptome increased significantly (from 32.5% to 45.6% for the nuclei and from 49.4% to 58.8% for the cells; Data S2). We obtained 8517 nuclei and 8564 cells that passed QC thresholds (on gene counts and mitochondrial reads, see Methods). The median number of genes detected was 1351 and 1506 for nuclei and cells, respectively. After batch effect correction, we combined these cells and nuclei into a single dataset containing 17,081 high quality transcriptomes.

Clustering parameters such as the number of principal components used, k-nearest neighbor and cluster resolution resulted in different numbers of clusters and cluster sizes. Since previous knowledge on the expected number of cell types or their molecular markers was virtually nonexistent, we assigned a stability value to each cluster in order to detect meaningful cell types (Fig. S2b)^29^. By subsampling and reclustering the dataset, we identified the optimal clustering parameters that resulted in the highest number of stable clusters. This resulted in 42 distinct stable clusters, which we presume related to bona fide cell types (Fig. 1e). Almost all the clusters contained data points from both cells and nuclei (Fig S2c). To allow further exploration of this atlas by the community, we made it available as a portal in SCope (https://scope.aertslab.org/#/Octopus_Brain/). In the following sections, we will describe several of these clusters in more detail, based on their spatial localization and/or expression of marker genes.

### Cluster annotation based on neurotransmitter and peptide expression

The majority of cells present in the octopus brain were neurons (89% *elav+*, 83% *onecut+*, Fig. S3a, Fig. 1f). A hierarchical clustering based on the transcriptomes of all stable clusters resulted in three main branches: neuronal, glial and other (hemocytes and endothelial cells; Fig. 1f). Several neuron types strongly exhibited a particular neurotransmitter or a peptidergic phenotype and were annotated accordingly, making use of gene homologs of fly and/or mouse (Fig. 2a). The paralarval brain was mostly glutamatergic (64% *vglut+*) and cholinergic (29% *vacht+*), but we also found four prominent dopaminergic clusters (27% of all cells are *th+*) (Fig. 2b). We observed that different cholinergic (ACH) and dopaminergic (DOP) neuronal clusters group together (fig 1f), which suggests a common origin and/or a common transcriptional program. On the t-SNE plot, a large central cluster of neurons was visible that we could not assign to a stable cluster. These cells were of high quality and could be divided in a cholinergic and glutamatergic population (Fig. 2b, fig S2d). The observation of such a central unstable cluster in the t-SNE was similar to what was seen in the *Drosophila* brain atlas^33^. A large set of neurons could not be clustered into distinct cell types in the fly brain, either, which may point to a large number of neuronal subtypes each with a small number of cells. Conversely, larger stable clusters likely represented more prevalent cell types.

**Fig. 2.**
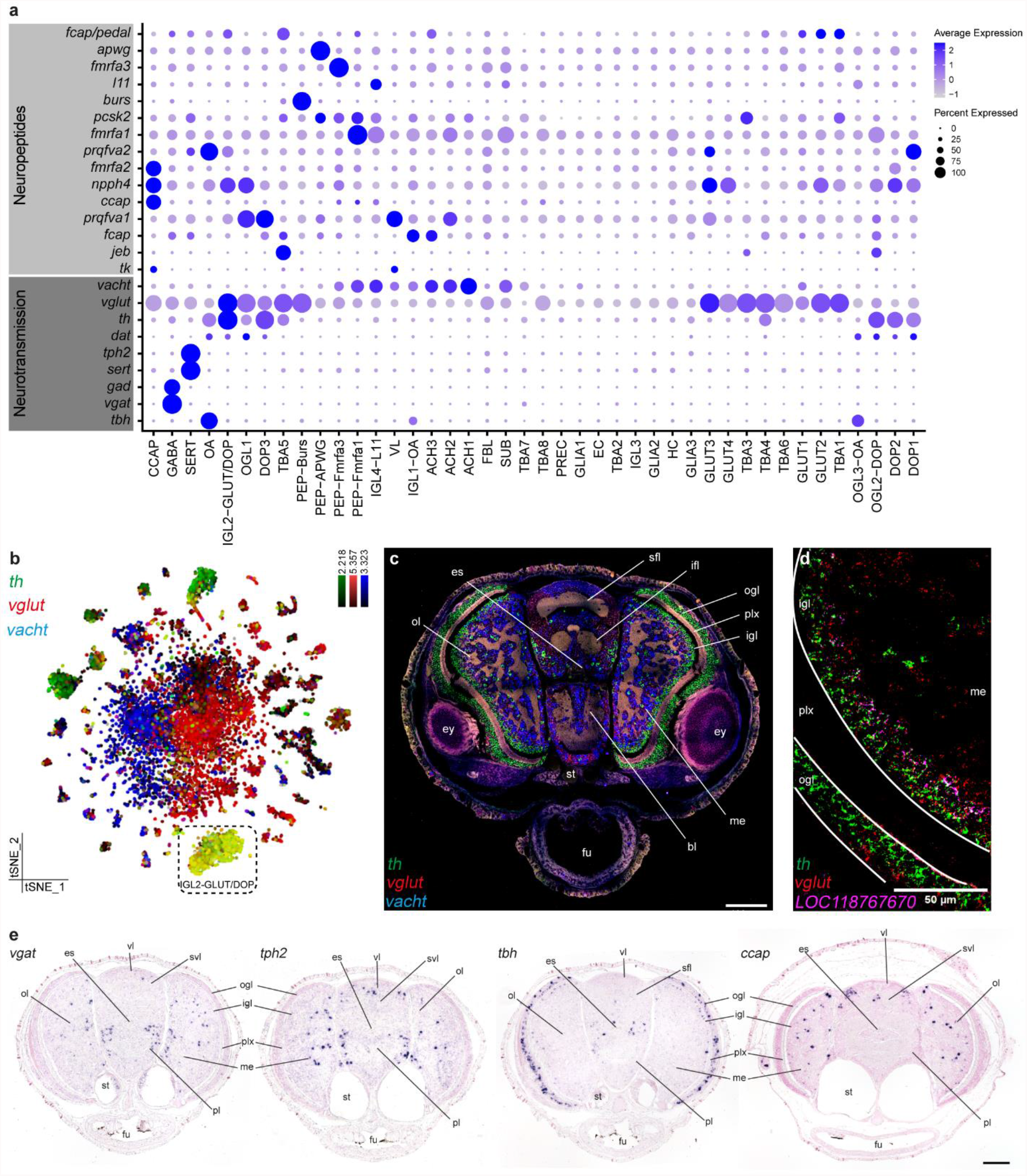
Neurotransmitters and peptides. **a** Dotplot for main neuropeptides and genes involved in the synthesis and transport of neurotransmitters. **b** Expression of *th, vglut* and *vacht* is visualized on a t-SNE plot. *Th* (tyrosine hydroxylase) is shown in green, *vglut* (vesicular glutamate transporter) in red and *vacht* (vesicular acetylcholine transporter) in blue. **c** Multiplexed *in situ* hybridization chain reaction (HCR) for *th, vglut* and *vacht*. **d** Co-expression of *th* and *vglut* in the inner granular layer of the optic lobe, together with the cluster specific marker for IGL2-GLUT/DOP; *LOC118767670*. **e** *In situ* hybridization for *vgat* (vesicular GABA transporter, GABAergic neurons), *tph2* (tryptophan hydroxylase 2, serotonergic neurons), *tbh* (Tyramine β-hydroxylase, octopaminergic neurons) and the neuropeptide *ccap* (Crustacean cardioactive peptide). Scale bars represent 100 µm for the overview images. bl, brachial lobe; es, esophagus; ey, eye; fu, funnel; igl, inner granular layer; ifl, inferior frontal lobe; me, medulla; ogl, outer granular layer; ol, optic lobe; pl, pedal lobe; plx, plexiform layer; sfl, superior frontal lobe; st, statocysts; svl, subvertical lobe; vl, vertical lobe.

In order to spatially locate cell types within the brain, we proceeded with *in situ* hybridization for highly expressed genes related to neurotransmitter synthesis or transport and genes encoding peptides (Fig. 2c-e). We identified a common dual-transmitter cell type, which is both dopaminergic and glutamatergic (Fig. 2b,d). *In situ* HCR showed that this cell type was prevalent in the inner granular layer (igl) of the optic lobe (IGL2-GLUT/DOP).

GABAergic (*gad+, vgat+*; GABA) and serotonergic (*sert+, tph2+*; SERT) neurons comprised smaller populations that were located throughout the medulla of the optic lobe and the central brain (Fig. 2e). Three octopaminergic cell types (OA) expressed the synthesizing enzyme *tbh* and could be distinguished based on their spatial localization (Fig. 2e); outer granular layer (OGL3-OA), inner granular layer (IGL1-OA) and central brain (OA). We could not identify any tyraminergic, glycinergic or (nor)adrenergic neurons. The majority of neurons expressed one or more neuropeptides, in addition to a neurotransmitter (e.g., OA; *tbh* and *prqfva2*). In contrast, some clusters did not have a clear neurotransmitter phenotype but did express a prominent neuropeptide, for instance *fmrfa3* (PEP-Fmrfa3) and *ccap* (CCAP) (Fig. 2e).

We also identified a cholinergic cell type (SUB) that was dedicated to the subesophageal mass. These neurons (Fig. 2c, Fig. S4) produced the neuropeptide *l11* and *shh* and were organized in groups of large cells within the sub. This cell type appeared intercalated with glutamatergic neurons (Fig. 2c).

### Molecular lamination within the deep retina

The optic lobe already contained a large cellular diversity at hatching. We further investigated whether neuronal subtypes were spatially confined or distributed by mapping subtype-specific marker genes. We found three distinct cell types within the outer granular layer (ogl) of the optic lobe (Fig. 3a,c,e,g, OGL1, *ppp1+*; OGL2-DOP, *jeb+*; OGL3-OA, *pcdh24+*). The majority of cells in the ogl were small dopaminergic neurons (OGL2-DOP). Cells in the OGL2-DOP cluster expressed *jeb* and *dscam* at different levels, which we confirmed using *in situ* hybridization (Fig. S5). *dscam*+ cells were located more towards the interior of the layer while *jeb*+ cells were positioned more externally (Fig. S5). Cell bodies of a second cell type (OGL1) seemed slightly larger than the dopaminergic cells and were mainly glutamatergic but also synthetized some dopamine. Lastly, the largest cell bodies we identified were octopaminergic (OGL3-OA). These octopaminergic neurons were a lot less prevalent than OGL1 and OGL2-DOP.

**Fig. 3.**
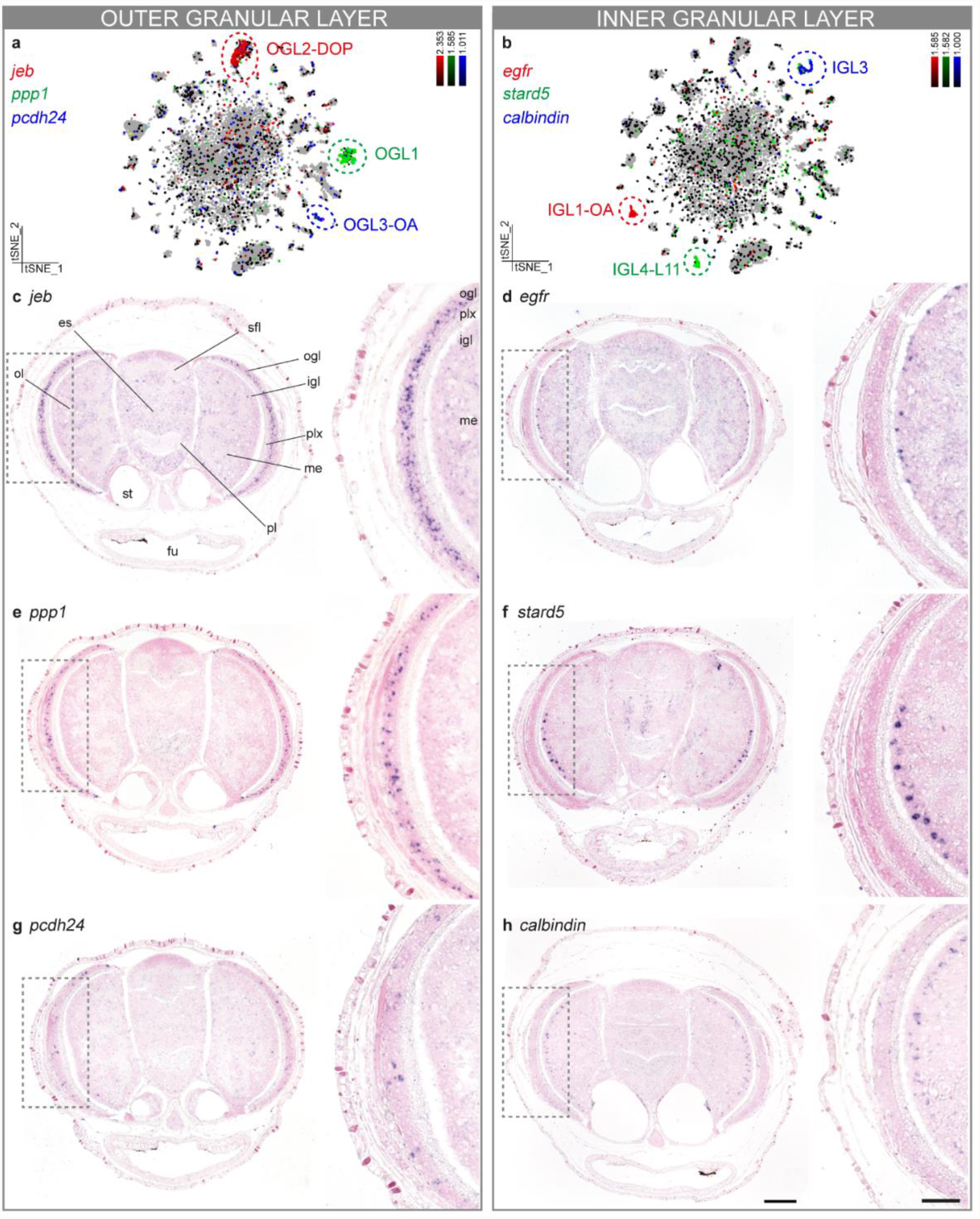
Optic lobe cell type diversity. **a** t-SNE representation of three different cell types (OGL1, OGL2-DOP, OGL3-OA) of the outer granular layer. Marker genes for these three populations are plotted. *In situ* hybridization for these marker genes are shown in c, e, g. **b** t-SNE representation of three cell types of the inner granular layer (IGL1-OA, IGL3 and IGL4-L11). Marker genes for these three populations are plotted. *In situ* hybridization for these marker genes are shown in d, f, h. **c** Neuropeptide *jeb* is expressed throughout the ogl in dopaminergic neurons. **d** Epidermal growth factor receptor (*egfr*) is expressed in octopaminergic neurons in the most outer region of the igl. **e** Protein phosphatase 1 (*ppp1*) is expressed within the ogl, although present in fewer cells than *jeb*. **f** StAR Related Lipid Transfer Domain containing 5 (*stard5*) is expressed within the igl, more interiorly than the *egfr+* cell type. **g** *pcdh24* is expressed in octopaminergic neurons present in the ogl. **h** *calbindin* is expressed in the most interior side of the igl. Magnified regions are annotated with a grey box. Scale bars for the overview images represent 100 µm and for the magnifications 50 µm. es, esophagus; fu, funnel; igl, inner granular layer; me, medulla; ogl, outer granular layer; ol, optic lobe; pl, pedal lobe; plx, plexiform layer; sfl, superior frontal lobe; st, statocysts.

Furthermore, we observed multiple cell types within the inner granular layer (igl) of the optic lobe (Fig. 3b,d,f,h). Large *egfr*+ cells (IGL1-OA) were located externally, next to the plexiform layer, while *stard5+* cells (IGL4-L11) and *calbindin+* cells (IGL3) were organized in layers more towards the medulla. Intriguingly, IGL3 cells did not synthesize any prominent neurotransmitter or neuropeptide. A fourth igl population, marked by the uncharacterized gene *LOC118767670* (Fig. 2d), was both glutamatergic and dopaminergic (IGL2-GLUT/DOP). The laminated appearance of molecularly different cell types revealed an additional subdivision within the so-called “deep-retina” of octopus.

### Cross-species cell type comparisons

In order to identify and annotate evolutionary conserved cell types, we performed comparisons between octopus, fly^33^ and mouse^32^ brain single-cell data sets using the SAMap algorithm^31^ (Fig. 4a). Based on cross-species cell type mappings, we found that the octopus GLIA1 subtype is molecularly similar to fly ensheathing glia and mouse astrocytes, and GLIA3 to mouse telencephalic astrocytes (Fig. 4a). Based on this mapping we could also identify a conserved glial gene expression signature that is shared between these three species (Fig. 4b). Only around 10% of all cells in the octopus paralarval brain were identified as glia (*gs2*+), see Fig. 4c. Both *gs2* and *apolpp* were highly expressed in all glial populations. In line with their function, we found that glial cells are located in the neuropil of the octopus brain (Fig. 4d-g). We then examined the expression of *apolpp* with *in situ* HCR at a higher magnification and were able to identify glial cells with multiple processes (Fig. 4e). Most *apolpp+* cells were located within the neuropil near the axons of the cells from the perikaryal layer, although some glial cells were infiltrating the cortex and were located between the neuronal cell bodies (Fig. 4f). We observed high expression of several invertebrate glial markers such as *CG6216, notch* and *eaat1* but no orthologues could be identified for genes used to discriminate between glial subtypes in flies (*indy, wrapper* and *alrm*)^33,36^. At least three distinct glial subtypes were identified within this dataset (GLIA1,2,3), suggesting there is also functional diversification within octopus glial cells. GLIA1 highly expressed *gat1* and *CG6126* while GLIA2 was characterized by *hhex* expression. IGL2-GLUT/DOP was a prominent cell type within the octopus visual system and had a similar molecular profile to fly T1 neurons (e.g. *eaat1* and *gilt1*) (Fig.4a). It remains to be investigated whether IGL2-GLUT/DOP neurons are the amacrine cells that provide feedback from the igl to the plexiform layer, similar to the fly T1 neurons from the medulla to the lamina.

**Fig. 4.**
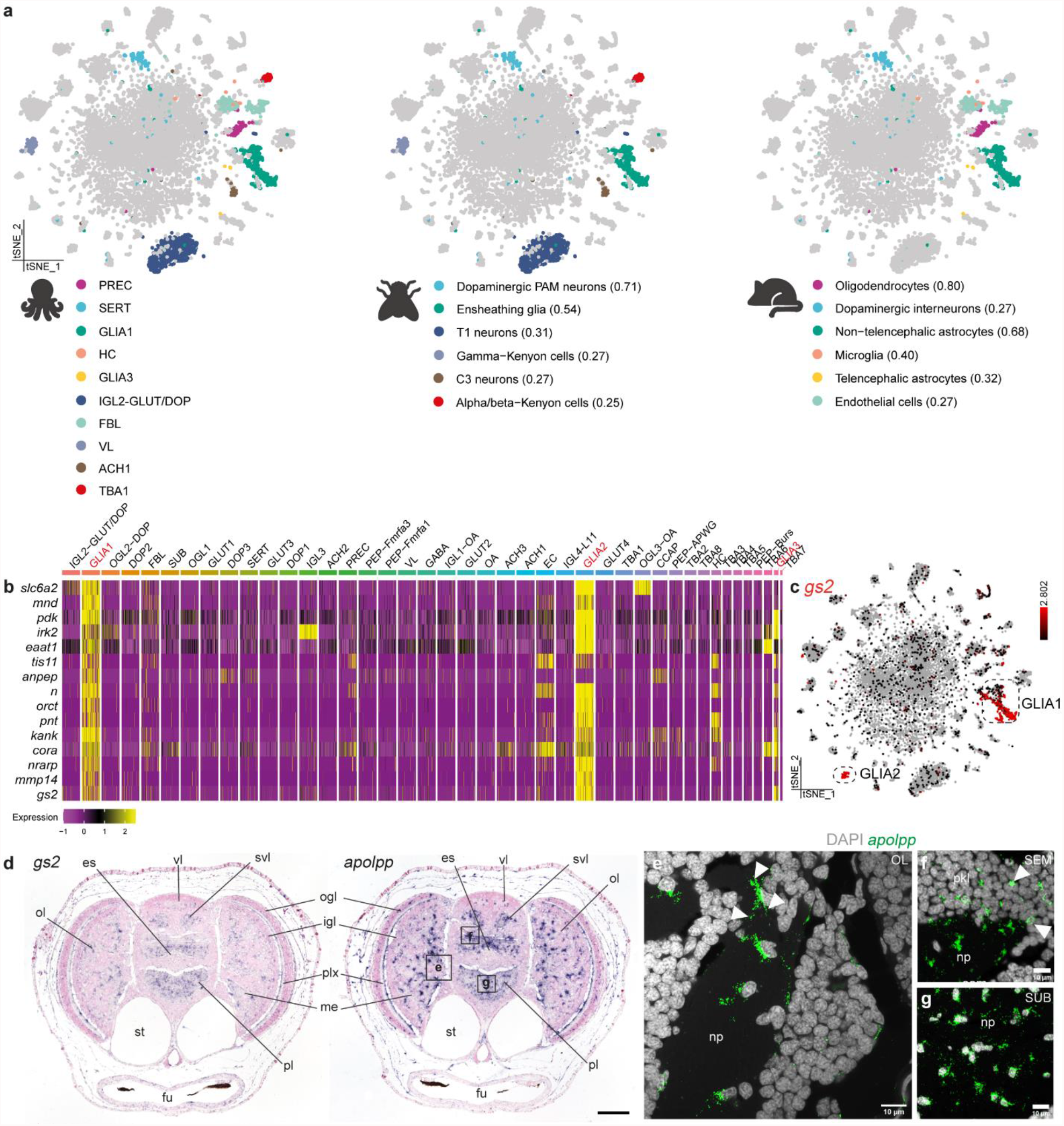
Cross species cell type comparisons identify a glial gene expression signature. **a** Cell type mappings between octopus, fly and mouse are represented on the t-SNE plot. Mappings are color coded and alignment scores are shown between brackets. **b** Heatmap of the top 15 genes (filtered on specificity and fold change) from the mapping between octopus glia 1, fly ensheathing glia and non-telencephalic astrocytes in the mouse brain. Glial populations are highlighted in red. **c** t-SNE representation of two main glial populations in octopus based on the expression of *gs2* (Glutamine synthetase 2). **d** *In situ* hybridization of *gs2* and *apolpp* (Apolipoprotein). Scale bar represents 100 µm. Representative magnifications of different brain areas are annotated with black boxes. Fluorescent *in situ* hybridization for *apolpp* are shown in e, f and g. **e** Glial cells in the optic lobe. White arrows indicate multiple processes. **f** Glial cells in the supraesophageal mass. White arrows indicate infiltrating glia. **g** Glial cells in the subesophageal mass. es, esophagus; fu, funnel; igl, inner granular layer; me, medulla; np, neuropil; ogl, outer granular layer; pkl, perikaryal layer; ol, optic lobe; pl, pedal lobe; plx, plexiform layer; SEM, supraesophageal mass; st, statocysts; SUB, subesophageal mass; svl, subvertical lobe; vl, vertical lobe.

SAMap also found similarities between octopus serotonergic neurons, fly dopaminergic PAM neurons and dopaminergic interneurons in mice (Fig.4a). In addition, fly lamina feedback C3 map to the octopus cell type ACH1. These findings suggest that some clusters might deploy deeply conserved transcriptional programs across bilaterian evolution.

Another interesting observation from the SAMap comparison was the similarity between octopus vertical lobe (VL) cells and fly gamma Kenyon cells (Fig. 4a). The VL cells were identified based on the expression of *aristaless* (*arx*), *camkII* and *tmtc4* (Fig. 5a). The adult *O. vulgaris* vl has five gyri, which consist out of 25 million small amacrine interneurons and 65,000 large neurons^6^. Although Frösh (1971) found that the relative volume of the vl was a lot smaller in hatchlings, the composition of the vl in the hatchling brain was not described^19^. We showed that the hatchling vl possesses 3-2 gyri along the dorsoventral axis and is composed of densely packed nuclei (Fig. 1c, 5a-c). This differs from observations made in *O. bimaculoides*, where five gyri could be readily distinguished after hatching^37^. Similarly, for *O. vulgaris*, we could not distinguish the amacrine cells from the large efferent neurons based on nuclear size. However, based on the widespread expression of these marker genes (*arx, camkII, tmtc4*) throughout the vl (Fig. 5a) and *vacht* expression, we could identify the VL cells as the cholinergic amacrine cells described in the adult brain^38^. Gene ontology enrichment analysis for this cell type showed that cognition, learning, and learning or memory are the top three most enriched biological processes. This supports the function of the vl as the structure contributing to associative processing and learning and memory in the octopus brain^39^. Regarding the molecular profile of these cells (Fig. 5d), genes involved in long term potentiation and memory formation (e.g., *calmodulin, camkII, rut*) were highly expressed. Common marker genes identifying the mushroom body in the fly, such as Dunce (*dnc*) and Leo (*pka-c*), were also enriched within VL cells^40^. Certain transcription factors (i.e. *mef2, mblk, dsp1* and *zfhx4)* were present in both VL and Kenyon cells (logfc.threshold > 0.25), but there was no significant enrichment for typical Kenyon cell transcription factors such as *ey, fru and dati*^33,41^.

**Fig. 5.**
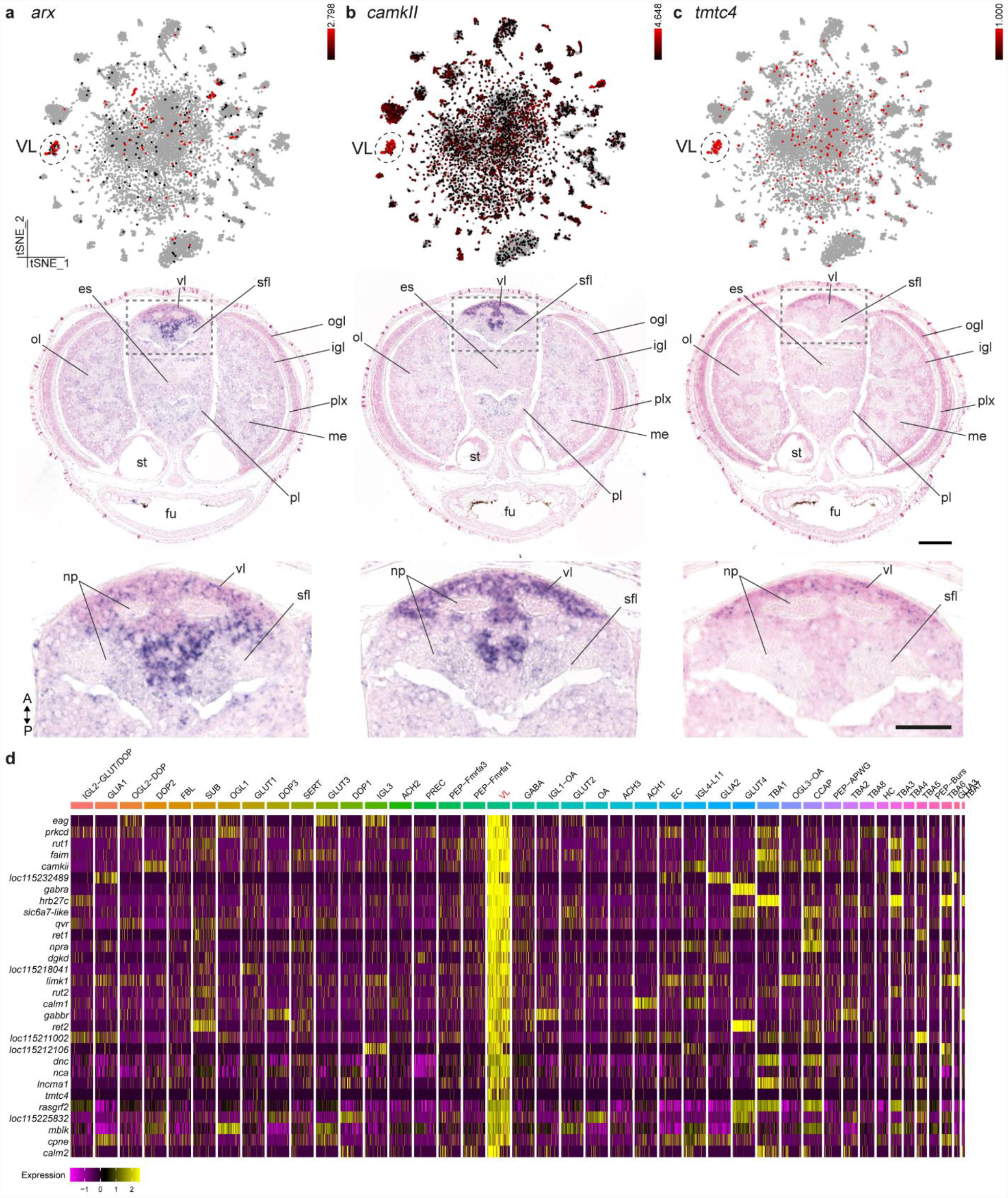
The molecular profile of the vertical lobe cells. **a** Aristaless (*arx*) expression is limited to the vertical (vl) and the superior frontal lobe (sfl). **b** Ca 2+ /Calmodulin-Dependent Protein Kinase II (*camkII*) expression can be observed mostly in the vl and less within the sfl. **c** Transmembrane O-Mannosyltransferase Targeting Cadherins 4 (*tmtc4*) is uniquely expressed within the most anterior part of the vl. t-SNE plots for *arx, camkII* and *tmtc4* are shown in a, b, c together with their respective *in situ* hybridization. **d** Top 30 genes with the highest fold change for the vertical lobe cells (VL). Scale bars are 100 µm for the overview images, 50 µm for the magnifications. es, esophagus; fu, funnel; igl, inner granular layer; np, neuropil; me, medulla; ogl, outer granular layer; ol, optic lobe; pl, pedal lobe; plx, plexiform layer; sfl, superior frontal lobe; st, statocysts; vl, vertical lobe.

### Non-neuronal cell types

In a previous study, we identified the lateral lips as the neurogenic niche outside of the developing octopus brain^18^.The lateral lips are anatomically very closely connected with the central brain through the anterior and posterior transition zones. We could retrieve limited expression of previously identified transcription factors (*ascl1* and *neurod*), which we assumed were lateral lip/transition zone cells (Fig. S6). These precursors (PREC) highly expressed markers related to pluripotency, embryonic stem cells and the npBAF complex. Genes such as *insm2, root* and a possible orthologue for *mki67* were highly expressed within the precursors. The majority of precursor cells were postmitotic (*neurod+*) but a smaller population were still progenitor-like (*ascl1+)* (Fig. S6b). Common markers for S and G2/M phase were highly expressed in this cluster (Fig. S6c). At this stage, we could only find a minor population of proliferating cells (PHH3+), within the remnants of the lateral lips but not in the brain (Fig. S6d). Interestingly, these precursors were found to be related to mouse oligodendrocytes (Fig. 4a). As invertebrates do not myelinate neurons, a myelinating cell type does not exist. The resemblance with mouse oligodendrocytes might point to a common ancestral cell type that has neural progenitor features. While this paralarval brain represents the end point of embryonic neurogenesis, a secondary phase of neurogenesis during a later stage is likely to occur.

Contrary to most invertebrates, the octopus has a closed circulatory system and a hemolymph-brain barrier^42–44^. At this developmental stage, we expected a certain degree of cerebral vasculature^45^. We found octopus endothelial cells (EC) that highly expressed conserved markers, more specifically *vegfr, hlx, meox2, troponin T* and *notch*. Furthermore, we identified a small population of hemocytes (HC) within the dataset (*vegf+, vwf+*). We also observed high *vegf* expression underneath the epidermis in a punctuate pattern (Fig. S7). The resemblance of the octopus hemocytes with mouse microglia, which are derived from the blood lineage, was not unexpected (Fig. 4a).

Fibroblast-like cells (FBL) were annotated based on their expression of collagens, troponin, tropomyosins and ribosomal genes. Intriguingly, hierarchical clustering of cell types illustrates similarities with precursor cells and neurons (Fig. 1f). Octopus fibroblasts were organized in a layer that surrounds the brain (Fig. S7). As this cell type produced extracellular matrix, it might contribute to forming the protective structure surrounding the central brain. Only half of the FBL expressed *troponin T* marking fully differentiated cells. Octopus FBL mapped to mouse endothelial cells, possibly owing to their common mesodermal origin (Fig. 4a).

### Homeobox genes are defining transcription factors for cell type identity

To investigate which transcription factors (TF) determine cell type identity, we calculated the tissue specificity index (tau) for all TF, which resulted in a ranked list. Then, we tested which TF family was the most cell type specific by performing a gene set enrichment analysis for the different TF families within the ranked list. We found that Homeobox TF are the most linked with cell type identity, followed by basic helix-loop-helix TF and ZnF, which massively expanded in coleoid cephalopods (Fig. 6a). Combinations of Homeobox TF do seem to be uniquely expressed in certain cell types (Fig. 6b). For example, we observed conserved expression of aristaless (*arx*) in amacrine interneurons within the vertical lobe (Fig. 5a) and *hlx* and *meox2* in endothelial cells. Moreover, we mapped *vsx2* to cells in the medulla and *prdl2*^46^ to the sub-vertical lobe (Fig. S8).

**Fig. 6.**
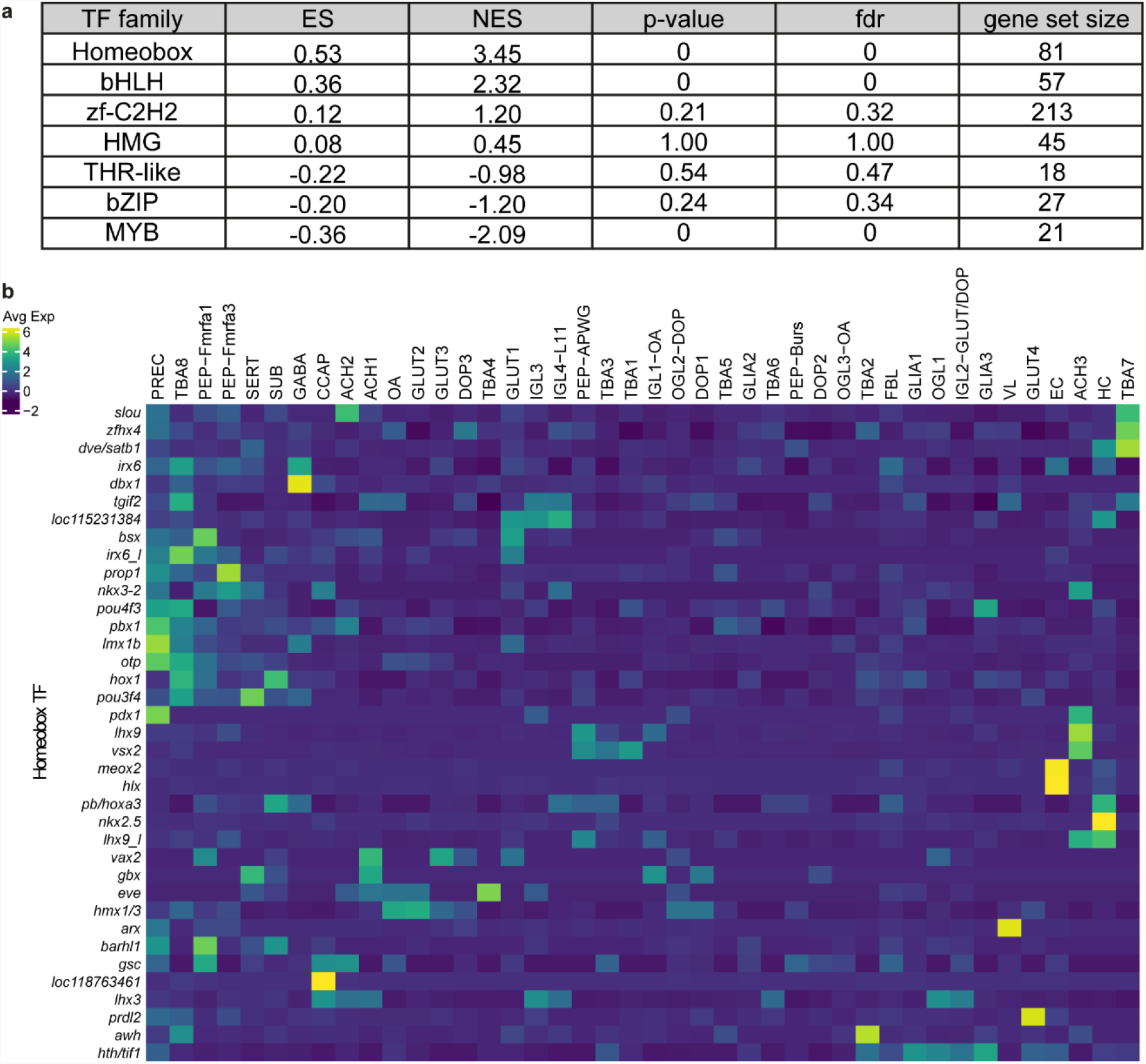
Cell type specificity and transcription factors. **a** Enrichment scores for the different TF within the ranked list based on tau. ES, Enrichment Score; NES, Normalized Enrichment Score; fdr, false-discovery rate. **b** Heatmap of highly variable Homeobox transcription factors, averaged per cell type and scaled.

### Genetic novelty drives cellular diversification

Our data showed a large diversity in brain cell types, which is expected in an animal with a rich cognitive behavioral pattern. Previous genomic studies indicated that coleoid cephalopods, including *O. bimaculoides* and *O. vulgaris*, specifically expanded certain gene families, leading to novel octopus genes^12,47^. We investigated whether these unique octopus genes might have driven the appearance of novel cell types. We hypothesized that recently expanded gene families, such as PCDH, ZnF, and GPCR, might convey the potential to diversify and develop octopus-specific cell types. For this purpose, we investigated whether genes of these families are enriched in certain cell types, which could be considered as a metric for novelty. In particular, PCDH were often annotated as marker genes (logfc.threshold > 0.25) for specific cell types. We found that some PCDH were ubiquitously expressed, while others were enriched in specific cell types (Fig. 7a). Important to note is that these PCDH were not homologous to vertebrate PCDH. *pcdh15* (Fig. 7b) was highly expressed within serotonergic neurons (SERT), whereas *pcdh24* (Fig. 7c) was enriched in a subset of octopaminergic neurons in the ogl (OGL3-OA). Although distinct subsets of GPCR and PCDH were highly expressed in specific neuronal cell types, the ZnF were enriched in the precursor cells (Fig. S9). This enrichment was statistically significant (p-adj<0.05) based on Fisher’s exact tests (bonferroni corrected) (see also Fig. S9b).

**Fig. 7.**
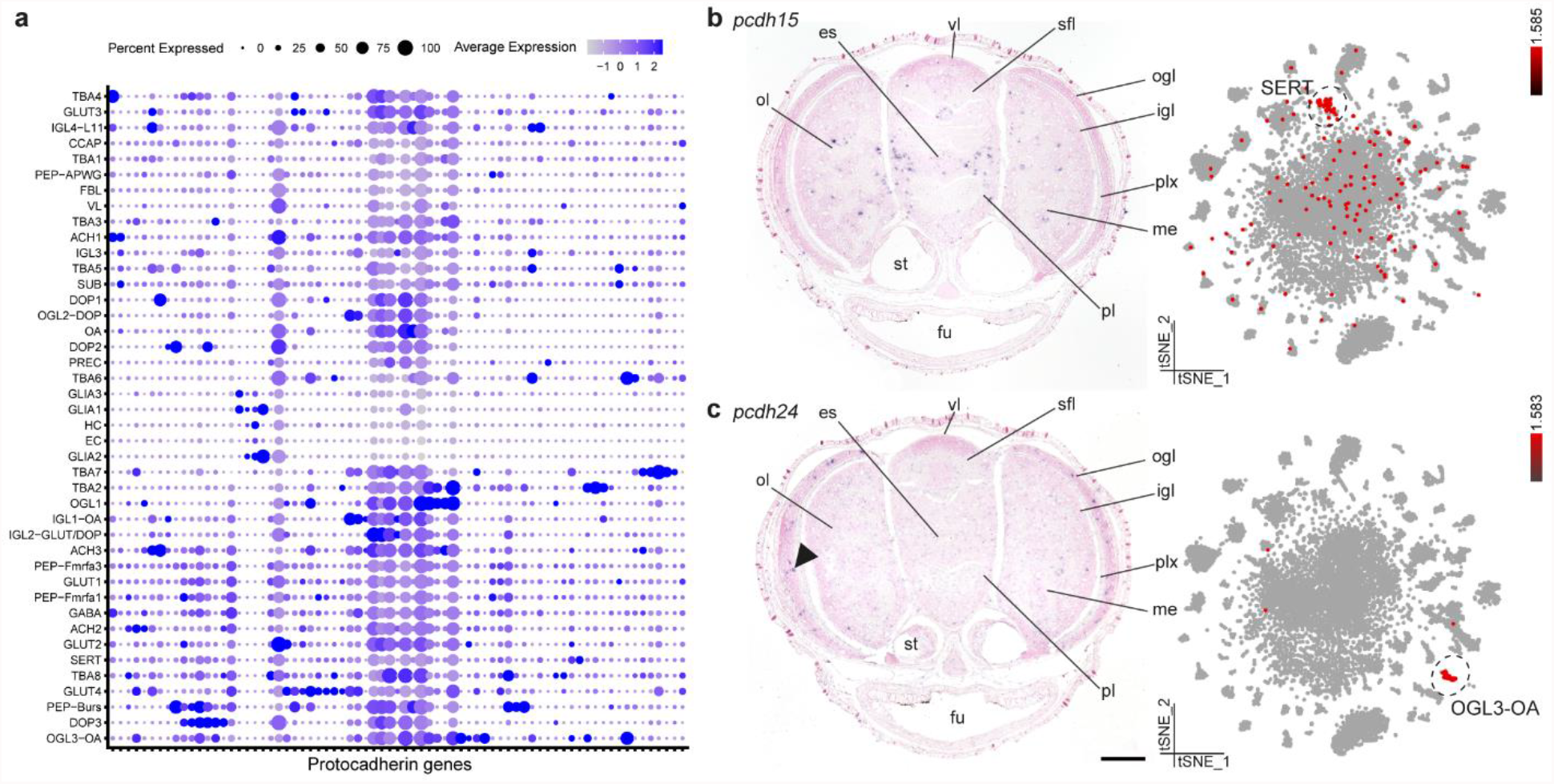
Protocadherin gene family expansion underlies cellular diversification. **a** Dotplot of highly variable protocadherin genes across all cell types. **b** Protocadherin 15 (*pcdh15*) is expressed within serotonergic neurons (SERT). **c** Protocadherin 24 (*pcdh24*) is expressed within the octopaminergic neurons localized in the outer granular layer (OGL3-OA), positive signal is indicated with a black arrow. Scale bars represent 100 µm. es, esophagus; fu, funnel; igl, inner granular layer; me, medulla; ogl, outer granular layer; ol, optic lobe; pl, pedal lobe; plx, plexiform layer; sfl, superior frontal lobe; st, statocysts; vl, vertical lobe.

## Discussion

Here, we report for the first time on cell type diversity in a mollusk. Although single-cell transcriptomic studies are gaining popularity, Lophotrochozoans have so far been understudied aside from *Platynereis dumerilii*^48^. In this study, we overcame several technical hurdles when applying novel technologies to a non-model organism such as *O. vulgaris*. Firstly, an evidence-guided approach to improve the genome annotation by using full-length RNA sequencing methods led to more complete gene models, including a better annotation of 3’UTRs and annotation of novel genes. This significantly improved mapping statistics and led to more reliable results and a higher number of estimated cells. Even in established model organisms such as zebrafish, a similar approach proved valuable and led to the identification of additional cell types^49^. We believe that this method and resource (Data S1) will aid other researchers in mapping bulk and scRNA-seq datasets. Secondly, cell type annotation without a priori knowledge on the number or molecular markers of cell types is not trivial. We propose that using cluster stability in order to obtain biologically relevant cell types^29^ is a reliable first way of cell type discovery. Lastly, by using a dual approach in sequencing both cells and nuclei, we have overcome any technical bias associated with each sequencing technology.

We were able to sequence around 17,000 single-cell expression profiles, which is roughly 9 percent of the total number of cells in the paralarval brain^20^. As a consequence, our data is unlikely to reveal every cell type present. Indeed, many neurons could not be assigned to stable clusters (Fig. 1e). Similar observations were made in adult fly brain scRNA-seq data^33,50^. We therefore hypothesize that the cell type diversity in these invertebrate brains is extensive and that the central cluster represents less prevalent cell types. Considering that this is a paralarval brain, of which the number of cells still needs to multiply a thousand-fold to reach adulthood^5,20^, the diversity of mature neuron types is impressive. Aside from a relatively small precursor population (∼1%), this brain seems fully differentiated. Interestingly, a prominent dual-transmitter cell type was identified (∼5% is both glutamatergic and dopaminergic). Similarly, in the larval fly brain 9% of neurons co-expressed markers for glutamatergic and aminergic neurons^36^, while this cell type was less prevalent in the adult fly brain^51^. These dual-transmitter neurons may play a role during development to refine the neural circuit and we postulate that this cell type might be unique to the optic lobe at this developmental stage.

The organization of the cell types within the inner and outer granular layer (igl and ogl) of the optic lobe is reminiscent of the very well characterized vertebrate retina^52,53^. We found that the ogl consists of at least three molecularly distinct cell types, which seem to be organized in sub-layers (Fig. S10). We identified numerous small cell bodies with a slight increase in size towards the exterior of the layer, which are most likely all amacrine neurons^53^. Although previous studies have identified eight different layers within the plexiform zone^53^, no lamination had been described so far within the igl. A laminated neuronal architecture coordinates visual information capture in vertebrates, in the layered retina, as well as visual information processing in invertebrates, in the optic lobe medulla in the fly. In these structures, layers are formed during development in a temporally controlled manner^54,55^. In a previous study, we have shown that the cells from the hatchling optic lobe originate from the lateral lips, which are spatially patterned^18^. Cells from the dorsal-anterior quadrant predominantly generate cells destined for the optic lobe cortex, whereas cells originating from the posterior lateral lip migrate towards the medulla. In addition to spatial patterning, future research is required to confirm whether these cells are also temporally patterned in order to generate the distinct layers within the optic lobe cortex.

While in mammals there are generally more glial cells in the brain than neurons, the opposite is true for most invertebrate species^56^. Considering that this is an unmyelinated and large central nervous system, cephalopods evolved alternative strategies to ensure conduction speed, e.g. the famous giant axon in squids^57^. Aside from myelin producing glia, wrapping glia that insulate axons can contribute to increased signaling speed^58^. Based on the location of glial cells within the octopus brain, we can differentiate between neuropil glia, presumably involved in axon wrapping, and infiltrating glia. The largest glial cluster identified (GLIA1) mainly expresses *gat*, which is also found in neuropil glia or astrocytes in the fly brain^36^. Whereas in the fly a distinctive glial type is identified at the borders of the neuropil, which is exclusively responsible for ensheathing the neuropil^59^, we could not identify a similar cell type based on the expression patterns of glial marker genes. The infiltrating glia likely provide support (both structural and metabolic) and might be involved in neuronal modulation as has been described for vertebrate astrocytes^60^, since they are in close proximity to the neuronal cell bodies. Whether this glial subtype correlates with GLIA2 cells is unknown at this point. SAMap analysis revealed a common glial gene expression signature between members of Lophotrochozoa, Ecdysozoa and chordates, suggesting that those genes reflect an ancestral bilaterian expression signature.

Conservation of gene expression profiles might point towards a common origin (out of an ancestral cell type) or a common function (by means of convergent evolution). In essence, SAMap does not make that distinction. Evolution of novel cell types from a common ancestor assumes the evolution or diversification of combinations of transcription factors or selectors^61,62^. Octopus vertical lobe cells were found to share expression of genes related to learning and memory in *Drosophila* Kenyon cells. The vertical lobe (vl) is considered to be the learning and memory center in octopus^38,39,63^. After ablation experiments of the vl, memory formation was found to be impaired^39^. Based on its ‘fan-out fan-in’ matrix-like synaptic network, its folded anatomy, small interneurons and the existence of LTP, this structure is possibly functionally homologous to a mushroom body^9^. Mushroom body-like structures have been identified in other lophotrochozoans such as *Platynereis* and a common origin has been suggested^64,65^. However, the overlapping expressed genes in octopus VL and *Drosophila* Kenyon cells did not contain many transcription factors specific to Kenyon cells. This suggest that different transcriptional programs have evolved to generate neurons involved in memory formation, and that the transcription factor code is more flexible than the underlying effector genes.

On the other hand, the VL cells highly expressed *arx*, which is a central transcription factor demarcating the early mushroom body in the annelid *Platynereis*^64^. Other transcription factors that have been commonly found in anterior brain structures in *Platynereis* and vertebrates, such as *pax6, emx2, lhx6, nkx2-1* or *dlx*, however, were not found to be expressed in the VL cells^66^. Similar to the *Drosophila* Kenyon cells, octopus VL cells are mainly cholinergic^38,67^. Future cell type atlases on a more diverse range of organisms might clarify whether these cell types share a common origin. The molecular blueprint of the VL cells could shed light on universal mechanisms regarding learning and memory. A direct link between *camkII* and memory storage has only recently been established in rats^68^. Similarly, octopus *camkII* signaling might also underlie the neuronal plasticity observed in the vl.

Transcription factors are considered major drivers of cell type identity. Recent studies in *C. elegans*^69^ and in fly motoneurons^70^ suggest that unique combinations of homeobox transcription factors are responsible for maintaining cell type identity. Our data show that that also in *O. vulgaris*, homeobox transcription factors are the most cell type specific. Together, our data suggest that the concept of a cell type determining homeobox code is translatable to organisms with increased neuronal cell type diversity such as *O. vulgaris*.

Expansion of ZnF, GPCR and PCDH gene families might have offered a way to develop novel cell types. Our data show that specific combinations of ZnF, GPCR and PCDH genes were expressed in particular clusters. By neo-functionalization of species-specific genes, unique cell types can arise. Indeed, novel genes have been found enriched in species-specific cell types^71^. The cell type-specific combinations of PCDH could have aided the development of the complex octopus nervous system^47^. Similar to non-clustered PCDH in vertebrates^72^, octopus PCDH combinations could provide a way to differentially sort out cell populations based on the adhesive character of PCDH. ZnF of the C2H2 type are amongst the most common DNA binding proteins in eukaryotes. ZnF were enriched in neuronal precursors, pointing to a potential role in cell fate specification and differentiation. This corroborates the finding that ZnF genes are more highly expressed during embryogenesis in *O. bimaculoides*^12^.

It remains an open question whether larger nervous systems also have more cell types or whether they have an increased cell number per cell type. Here we provide the first view on cell type diversity of a highly complex invertebrate brain, which we have only begun to explore. This dataset provides a starting point for comparative studies with the adult octopus brain, which might then yield informative answers linking brain complexity and cell type diversity. More cell numbers of certain cell types might increase the computational power of the brain, which could explain the higher cognitive function of the octopus brain.

## Supporting information

Description of Supplementary Data

## Data Availability

All single cell and nuclei data are available online at https://scope.aertslab.org/#/Octopus_Brain/. Scope allows for easy simultaneous visualization of the expression of three genes while toggling between different embeddings. Marker gene lists can be downloaded here for different clusterings (Seurat clustering and annotated clustering). Different metrics can be visualized such as the nCount, percent.mito, nFeature and whether these originated from cells or nuclei (batch). The scRNA-seq data and snRNA-seq data have been deposited in GEO under accession code GSE193622.

## Acknowledgements

The authors would like to thank Eduardo Almansa for his continuous support in providing us with octopus eggs and for sharing his expertise in octopus culture. Moreover, we want to thank Tania Aerts, Luca Masin and Ayana Rajagopal for critical discussions and Nikolai Hecker and Seppe De Winter for their help with bioinformatic analysis. We would also like to acknowledge our master thesis students Swell Sieben for help during dissections and Sofia Maccuro and Dries Janssen for cloning. All computational analyses were performed at the Vlaams Supercomputer Center (VSC). This work was supported by KU Leuven (C14/21/065 to E.S., ID-N/20/007 to E.S., S.A., C14/18/092 to S.A.), Stazione Zoologica Anton Dohrn (R.S., G.P., G.F.) and FWO (A.D.; SB/1S19517N and A.M.E.; FR/11D4120N).

## Author contributions

R.S., S.P., A.M.E. and A.D. performed the experiments. R.S., G.Z., G.H., K.S., A.M.E. and E.S. analyzed and interpreted the data. S.A., G.F. and E.S. designed and supervised the study. R.S. and E.S. wrote the original draft of the manuscript. All authors contributed to review and editing.

