## Supplementary material for "Cell type diversity in a developing octopus brain": Description of Supplementary Data

### Description of Additional Supplementary Files:

**Title: Supplementary Data 1**
**Description: 3'-extended genome annotation *(.gtf)*** *This gene transfer format file describes the location of the new and extended gene models within the genome.*

**Title: Supplementary Data 2**
**Description: Sequencing metrics. *(.xlsx)*** *Comparison of mapping statistics of the single cell and single nuclei data against the genome annotation on NCBI and the 3' extended genome annotation.*

**Title: Supplementary Data 3**
**Description: Functional genome annotation. *(.xlsx)*** *Additional functional genome annotation based on multiple databases (EggNog5, Blast and NCBI). The top blast hit against Drosophila melanogaster (Dm), Mus musculus (Mm) and Octopus bimaculoides (Ob) are shown.*

**Title: Supplementary Data 4**
**Description: Cluster annotation. *(.xlsx)*** *Sheet 1: Annotation of all the identified Seurat clusters. Summary of the SAMap results against fly and mouse datasets and individual stability values for each cluster are reported. Marker genes that were used for in situ hybridization are also described. Sheet 2: Gene identification numbers of octopus genes mentioned within this manuscript.*

**Title: Supplementary Data 5**
**Description: Marker gene lists. *(.xlsx)*** *Sheet 1: Differentially expressed genes for all 87 Seurat clusters are listed (logfc.threshold = 0.25). Sheet 2: Differentially expressed genes for the 42 stable clusters are listed (logfc.threshold = 0.25).*

**Title: Supplementary Data 6**
**Description: Probe sequences for *in situ* hybridizations. *(.xlsx)*** *Sheet 1: Overview of all probes used for colorimetric in situ hybridization experiments. Probe sequences and number of replicates are listed. Sheet 2: Probe sets used for hybridization chain reaction (HCRv3). The number of probe pairs and the amplifier details are described.*
