## Supplementary Figures for "Cell type diversity in a developing octopus brain"

**This PDF file includes:**

**Figs. S1 to S10**

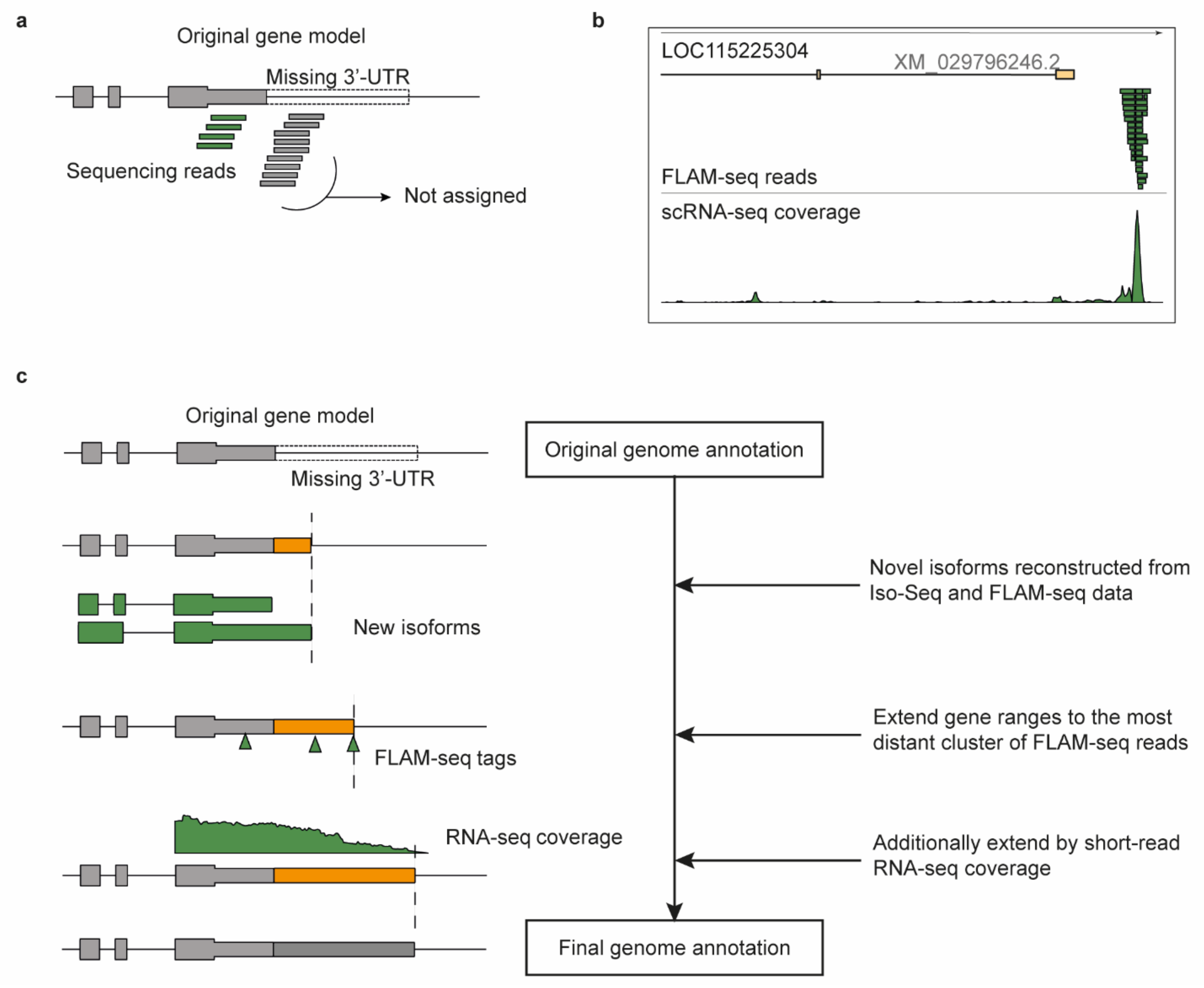


**Fig. S1 Extending the 3' gene annotation. a** Hypothetical gene with incomplete 3' UTR annotation. Sequencing reads are not counted. **b** *iqsec2* gene with incomplete 3' UTR annotation. The end of the 3' UTR is shown by the FLAM-seq reads, which coincides with read coverage of sc/snRNA sequencing data. **c** Schematic overview of the methods used to extend the gene annotation.

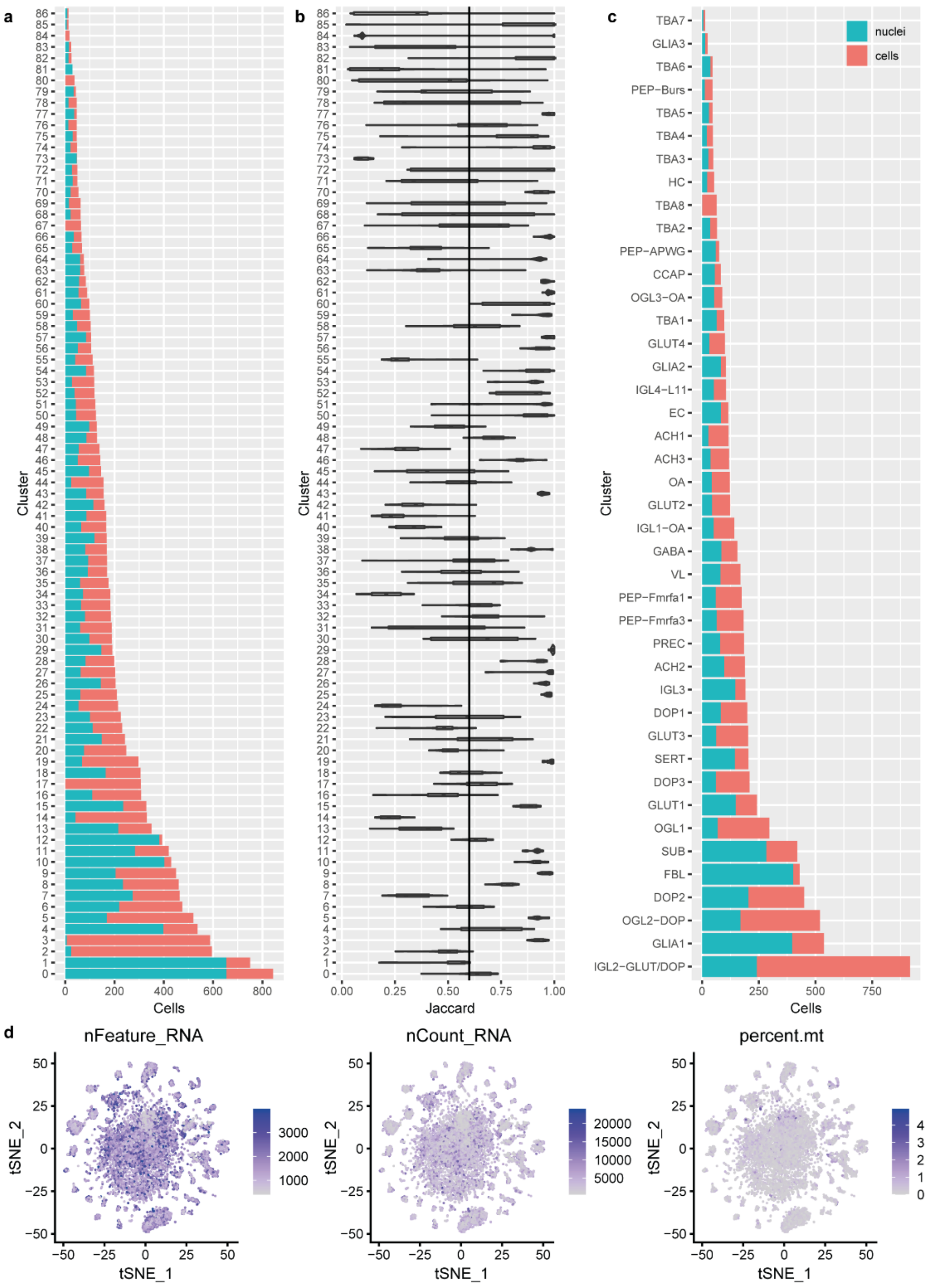


**Fig. S2 Data integration, clustering and quality control. a** Resulting clusters after data integration between cells and nuclei. The number of cells originating from either the cells or the nuclei are color coded. **b** Jaccard indices after subsampling and reclustering with parameters that result in the highest number of stable clusters (dims =150, k.param =10, resolution =2). Clusters with mean Jaccard indices above 0.6 are considered as stable clusters. **c** Resulting stable clusters after data integration between cells and nuclei. The number of cells originating from either the cells or the nuclei are color coded. **d** Multiple metrics for quality control of the resulting dataset are represented on the t-SNE plot.

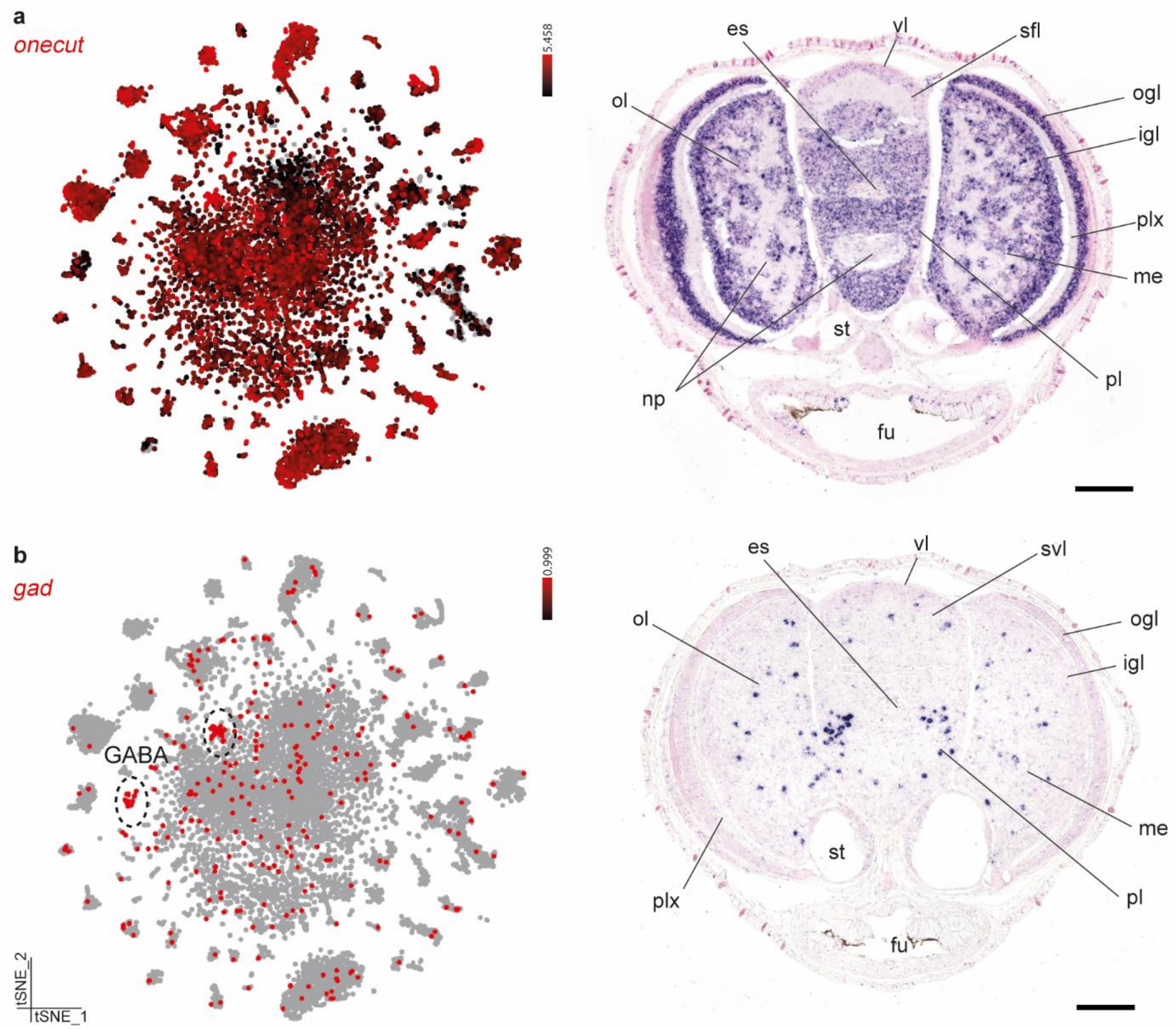


**Fig. S3 Neuronal markers. a** *onecut* expression is visualized on a t-SNE plot. *In situ* hybridization for *onecut* is shown on the right. Perikaryal layers are stained in purple and neuropil (np) is visible in pink. **b** t-SNE representation of GABAergic cells. Expression of Glutamate decarboxylase (*gad*) is shown in red. Scale bars represent 100 µm. es, esophagus; fu, funnel; igl, inner granular layer; me, medulla; np, neuropil; ogl, outer granular layer; ol, optic lobe; pl, pedal lobe; plx, plexiform layer; sfl, superior frontal lobe; st, statocysts; sfl, superior frontal lobe; svl, subvertical lobe; vl, vertical lobe.

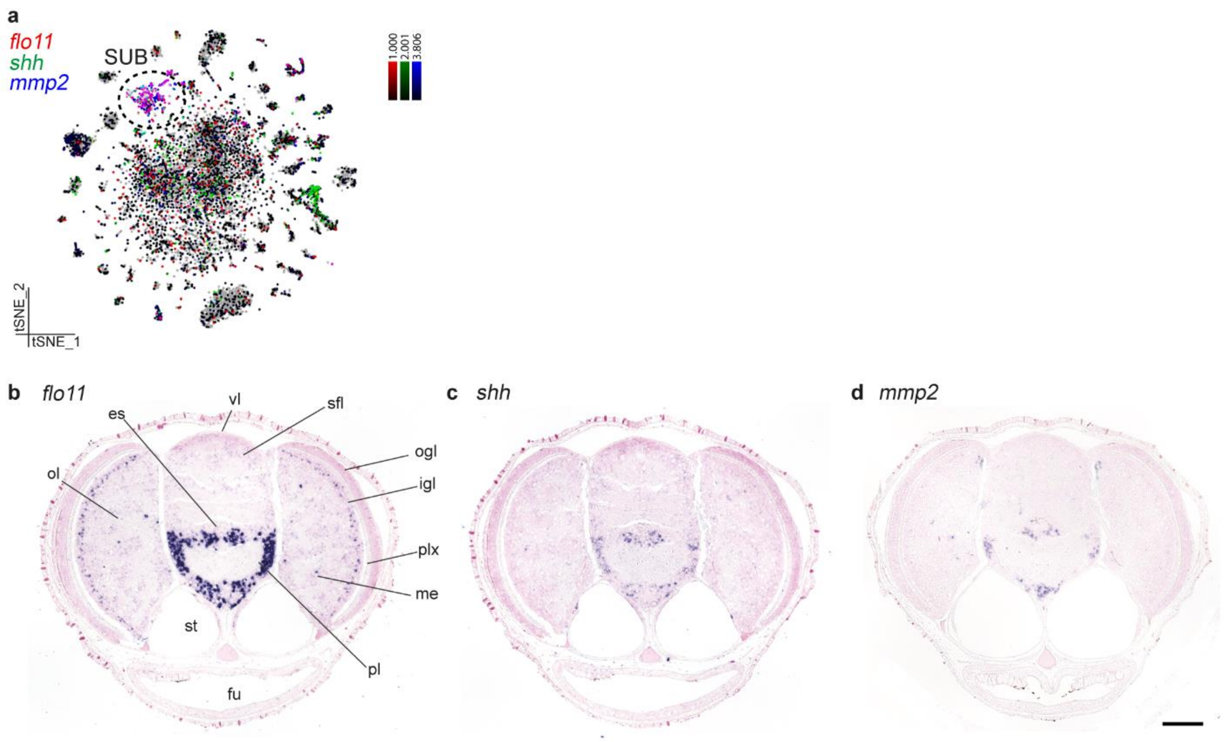


**Fig. S4 Sub-esophageal lobe cells. a** Cells colored by expression of *flo11* (red), *shh* (green) and *mmp2* (blue) visualized in a t-SNE plot. **b** Flocculation protein-like (*flo11*) is highly expressed within the sub-esophageal mass and in a smaller population in the igl. **c** Sonic hedgehog (*shh*) is expressed in the sub-esophageal mass and in subpopulation of the glial cells. **d** Matrix metallopeptidase 2 (*mmp2*) is expressed in the sub-esophageal mass. igl, inner granular layer; SUB, sub-esophageal mass. Scale bar represents 100 µm. es, esophagus; fu, funnel; igl, inner granular layer; me, medulla; ogl, outer granular layer; ol, optic lobe; pl, pedal lobe; plx, plexiform layer; sfl, superior frontal lobe; st, statocysts; vl, vertical lobe.

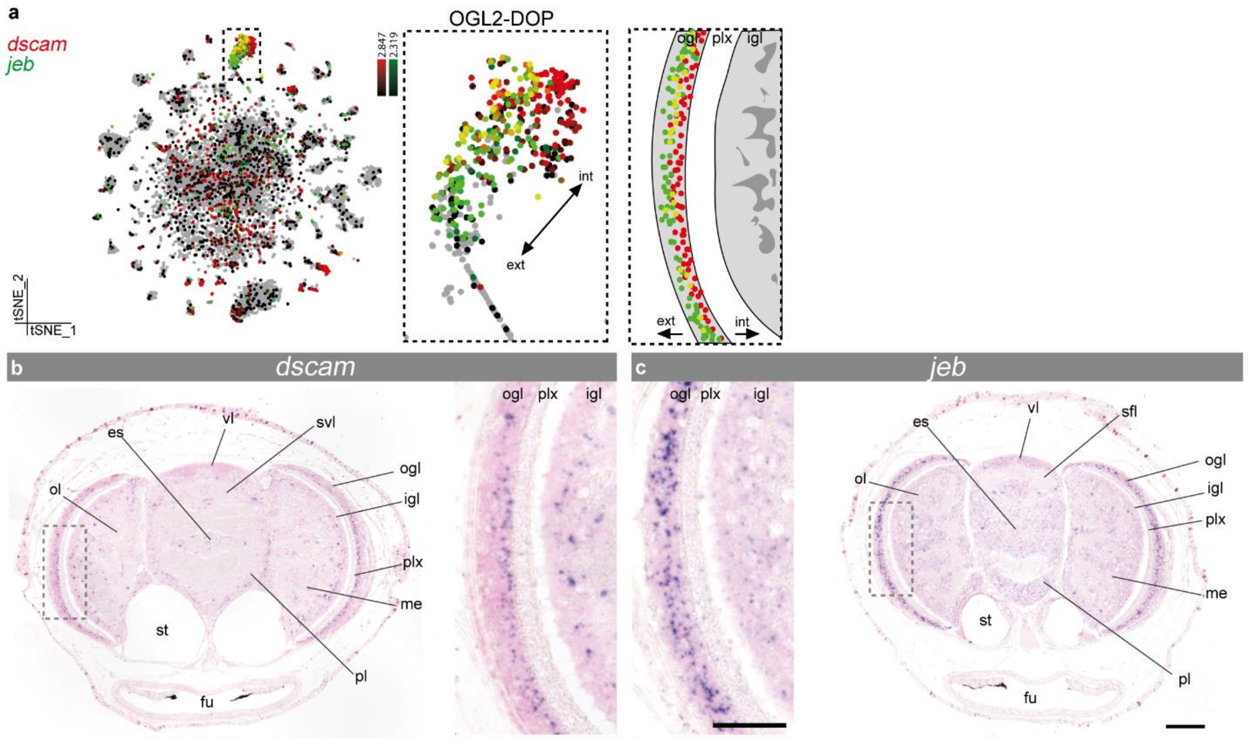


**Fig. S5 Dopaminergic cells in the outer granular layer. a** t-SNE representation of two different markers for the OGL2-DOP population. *dscam* (Down syndrome cell adhesion molecule) is shown in red and *jeb* (Jelly belly) in green. A magnification for OGL2-DOP is shown together with a color coded scheme for *jeb* and *dscam* expression. **b** *dscam* is expressed in small cells more towards the interior of the outer granular layer. **c** Neuropeptide *jeb* is expressed throughout the ogl in dopaminergic neurons. Magnified regions are annotated with a grey box. Scale bars for the overview images represent 100 µm and for the magnifications 50 µm. es, esophagus; fu, funnel; igl, inner granular layer; me, medulla; ogl, outer granular layer; ol, optic lobe; pl, pedal lobe; plx, plexiform layer; sfl, superior frontal lobe; svl, subvertical lobe; st, statocysts; vl, vertical lobe.

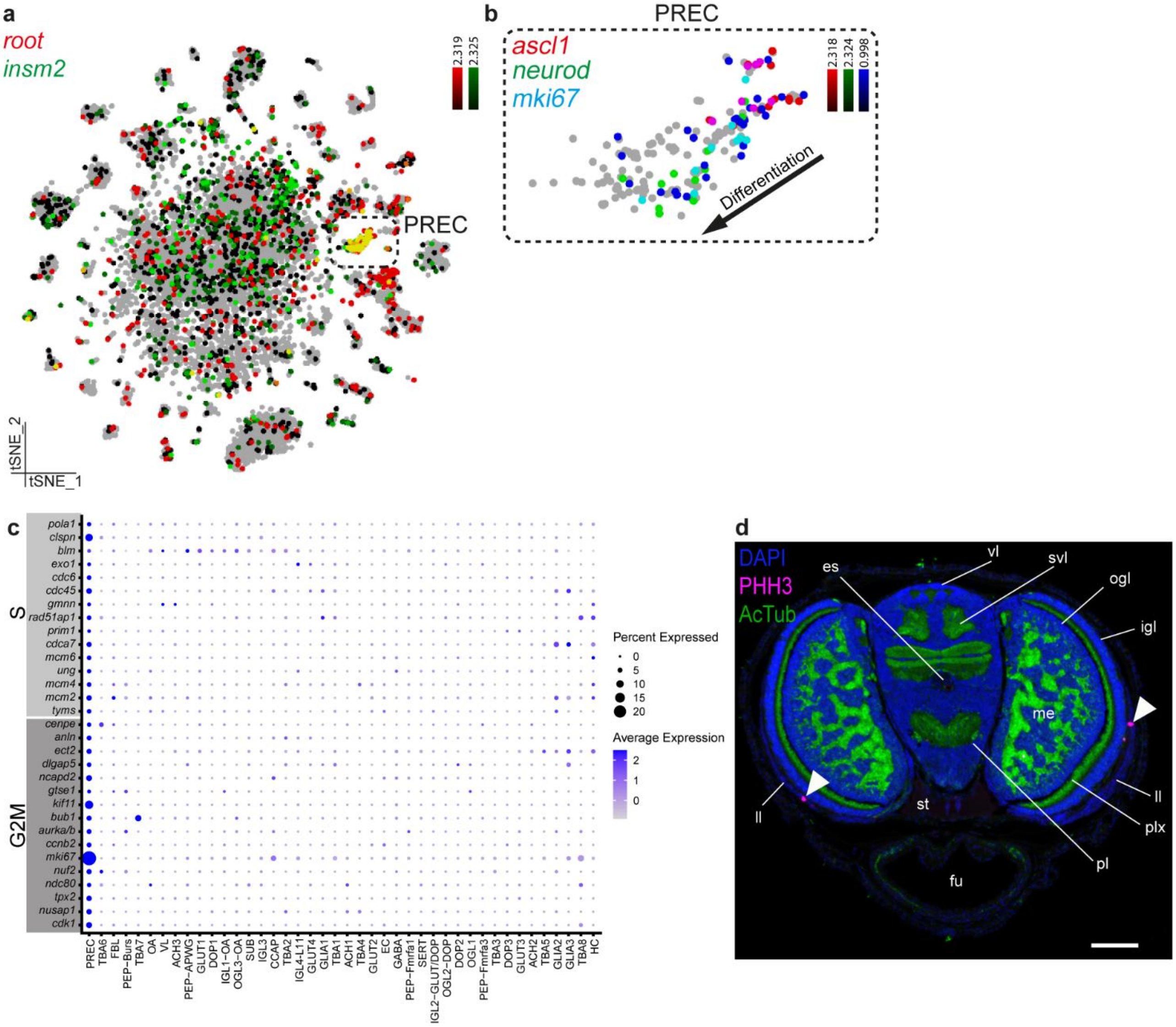


**Fig. S6 Precursor cells in one-day-old paralarvae. a** Expression of Rootletin (*root*) is shown in red and Insulinoma-Associated 2 transcriptional repressor (*insm2*) in green on the t-SNE plot. **b** Precursors express *ascl1*, *mki67* and *neurod*. A gradient in differentiation is visible. **c** Known markers for G2/M and S phases of the cell cycle are visualized on a dotplot. **d** Phospho-histone H3 (PHH3) staining on a one-day-old paralarvae, together with axonal marker acetylated tubulin (AcTub). Cells undergoing mitosis (late G2 and M-phase) are annotated with a white arrow. Scale bar represents 100 µm. es, esophagus; fu, funnel; igl, inner granular layer; ll, lateral lips; ogl, outer granular layer; pl, pedal lobe; plx, plexiform layer; st, statocysts; svl, subvertical lobe; vl, vertical lobe.

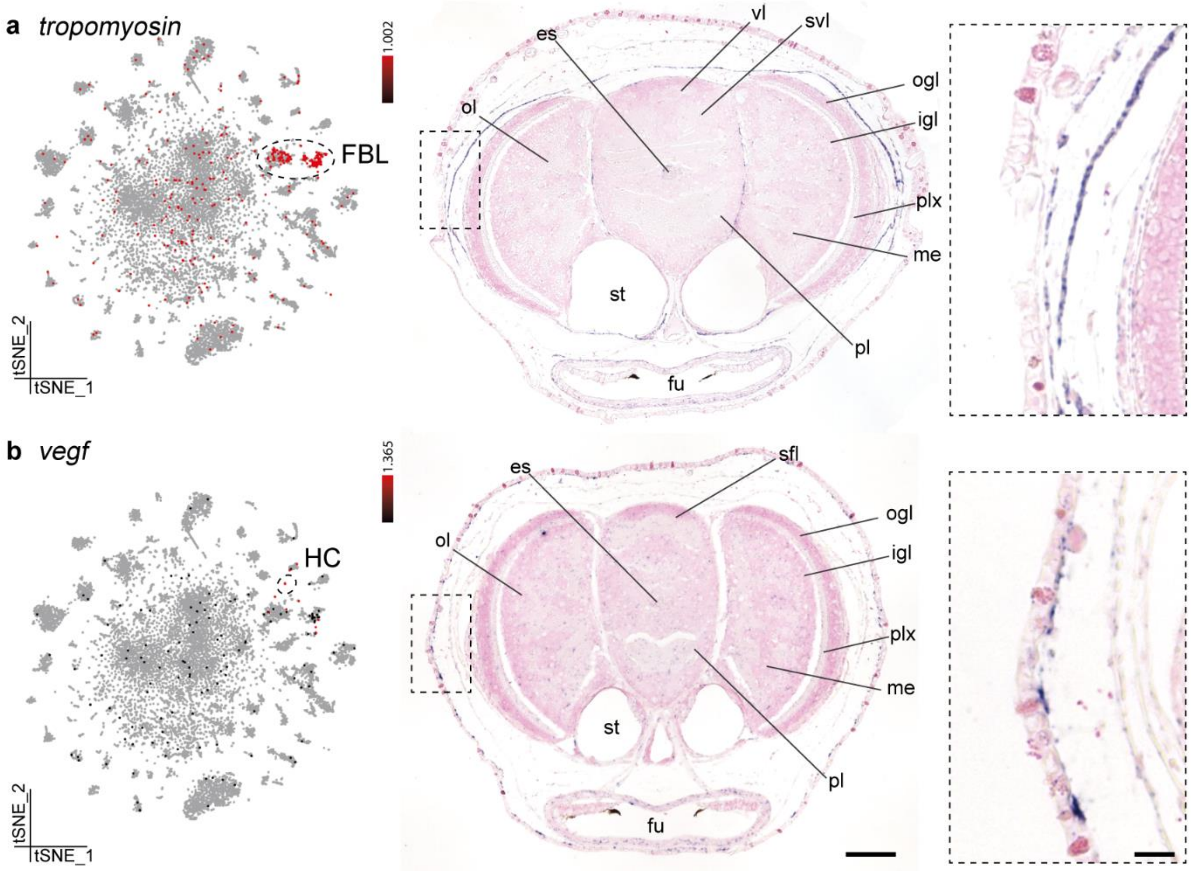


**Fig. S7 Non-neuronal cell types. a** *tropomyosin* is expressed within the fibroblasts (FBL). These cells form a layer that surrounds the brain. Expression can also be observed between the optic lobes and the central brain, and surrounding the statocysts. **b** *vegf* (vascular endothelial growth factor) is expressed within the hemocytes (HC). *vegf* expression is visible in a punctuate pattern underneath the epidermis. Scale bars for the overview images represent 100 µm and for the magnifications 50 µm. es, esophagus; fu, funnel; igl, inner granular layer; me, medulla; ogl, outer granular layer; ol, optic lobe; pl, pedal lobe; plx, plexiform layer; svl, subvertical lobe; sfl, superior frontal lobe; st, statocysts; vl, vertical lobe.

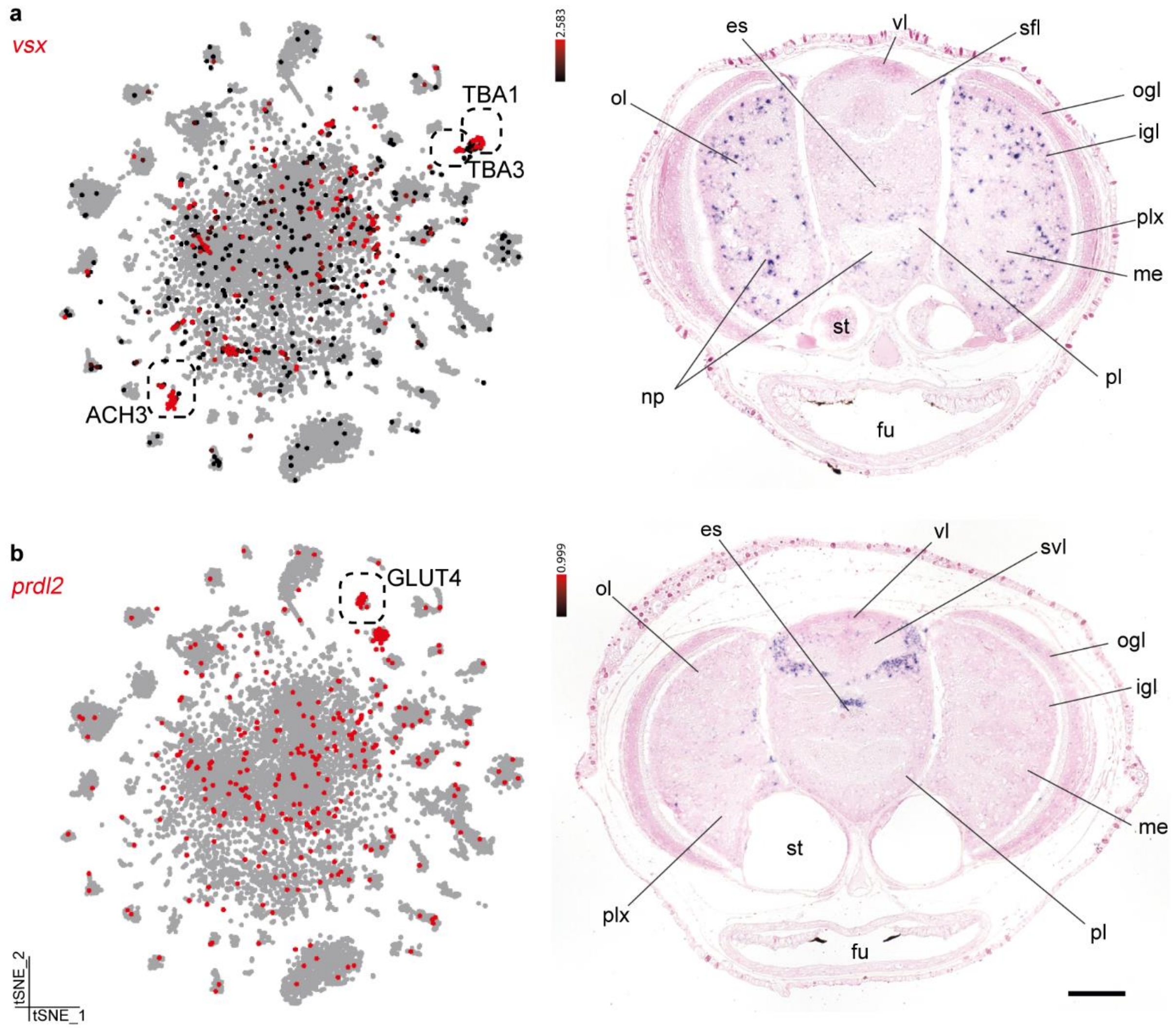


**Fig. S8 Gene expression profile of specific transcription factors. a** Visual system Homeobox transcription factor (*vsx*) expression is shown on the tSNE plot together with its *in situ* hybridization. *vsx* is expressed in TBA1, TBA3 and ACH3. These cells are distributed along the medulla of the optic lobes. Some expression is also visible within the sub-esophageal mass. **b** Paired homeobox protein-2-like (*prdl2*) expression is limited to the GLUT4 cluster and an unstable cluster (as shown on the tSNE on the left). With *in situ* hybridization we map these glutamatergic neurons to clusters of cells within the sub-vertical lobe. Scale bars represent 100 µm.  es, esophagus; fu, funnel; igl, inner granular layer; me, medulla; ogl, outer granular layer; ol, optic lobe; pl, pedal lobe; plx, plexiform layer; svl, subvertical lobe; sfl, superior frontal lobe; st, statocysts; svl, subvertical lobe; vl, vertical lobe.

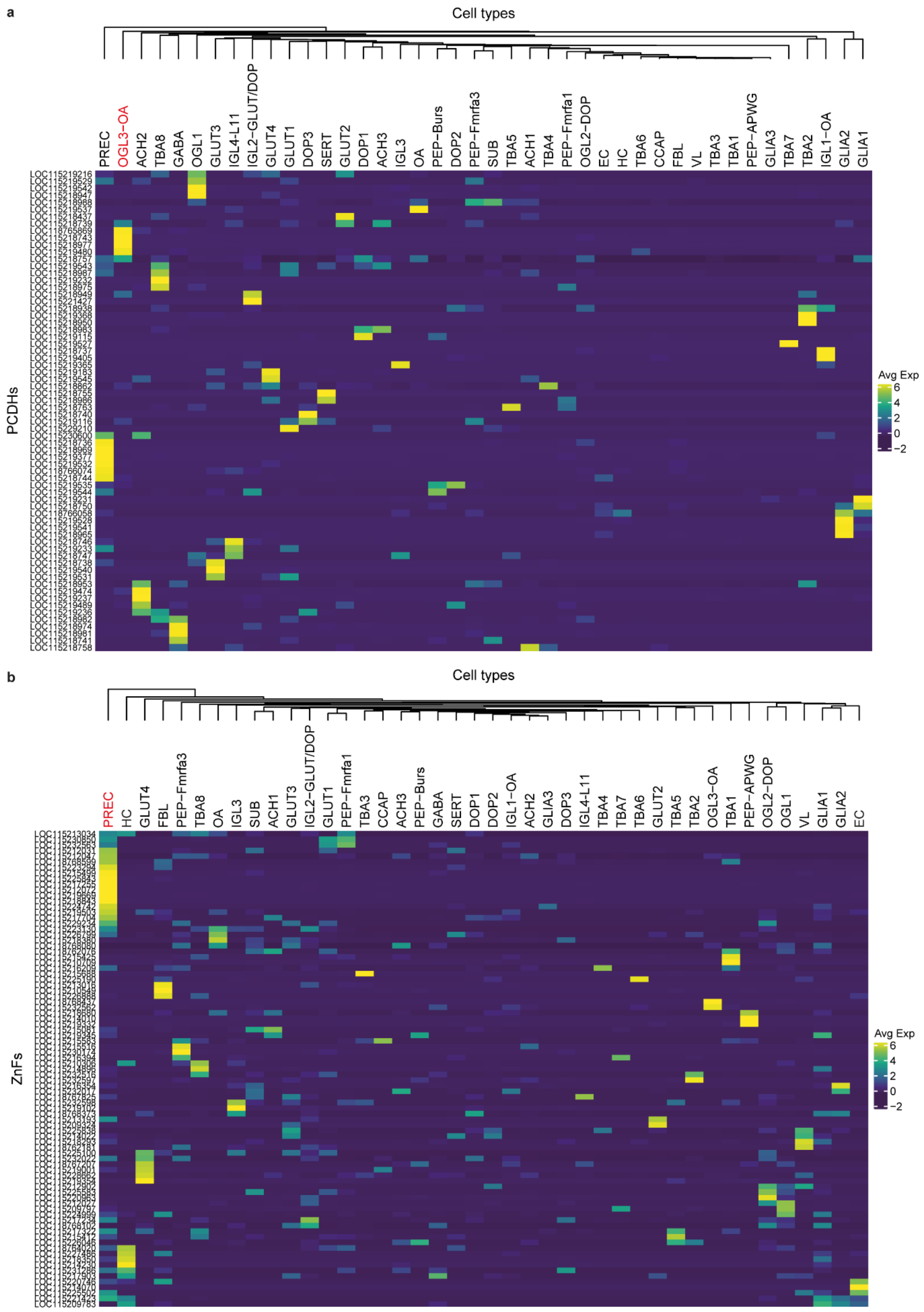
a
Cell types
PREC
OGL3−OA
ACH2
TBA8
GABA
OGL1
GLUT3
IGL4-L11
IGL2−GLUT/DOP
GLUT4
GLUT1
DOP3
SERT
GLUT2
DOP1
ACH3
IGL3
OA
PEP−Burs
DOP2
PEP−Fmrfa3
SUB
TBA5
ACH1
TBA4
PEP−Fmrfa1
OGL2−DOP
EC
HC
TBA6
CCAP
FBL
VL
TBA3
TBA1
PEP−APWG
GLIA3
TBA7
TBA2
IGL1−OA
GLIA2
GLIA1
PCDHs
Avg Exp
6
4
2
0
−2
b
Cell types
PREC
HC
GLUT4
FBL
PEP−Fmrfa3
TBA8
OA
IGL3
SUB
ACH1
GLUT3
IGL2−GLUT/DOP
GLUT1
PEP−Fmrfa1
TBA3
CCAP
ACH3
PEP−Burs
GABA
SERT
DOP1
DOP2
IGL1−OA
ACH2
GLIA3
DOP3
IGL4-L11
TBA4
TBA7
TBA6
GLUT2
TBA5
TBA2
OGL3−OA
TBA1
PEP−APWG
OGL2−DOP
OGL1
VL
GLIA1
GLIA2
EC
ZnFs
Avg Exp
6
4
2
0
−2

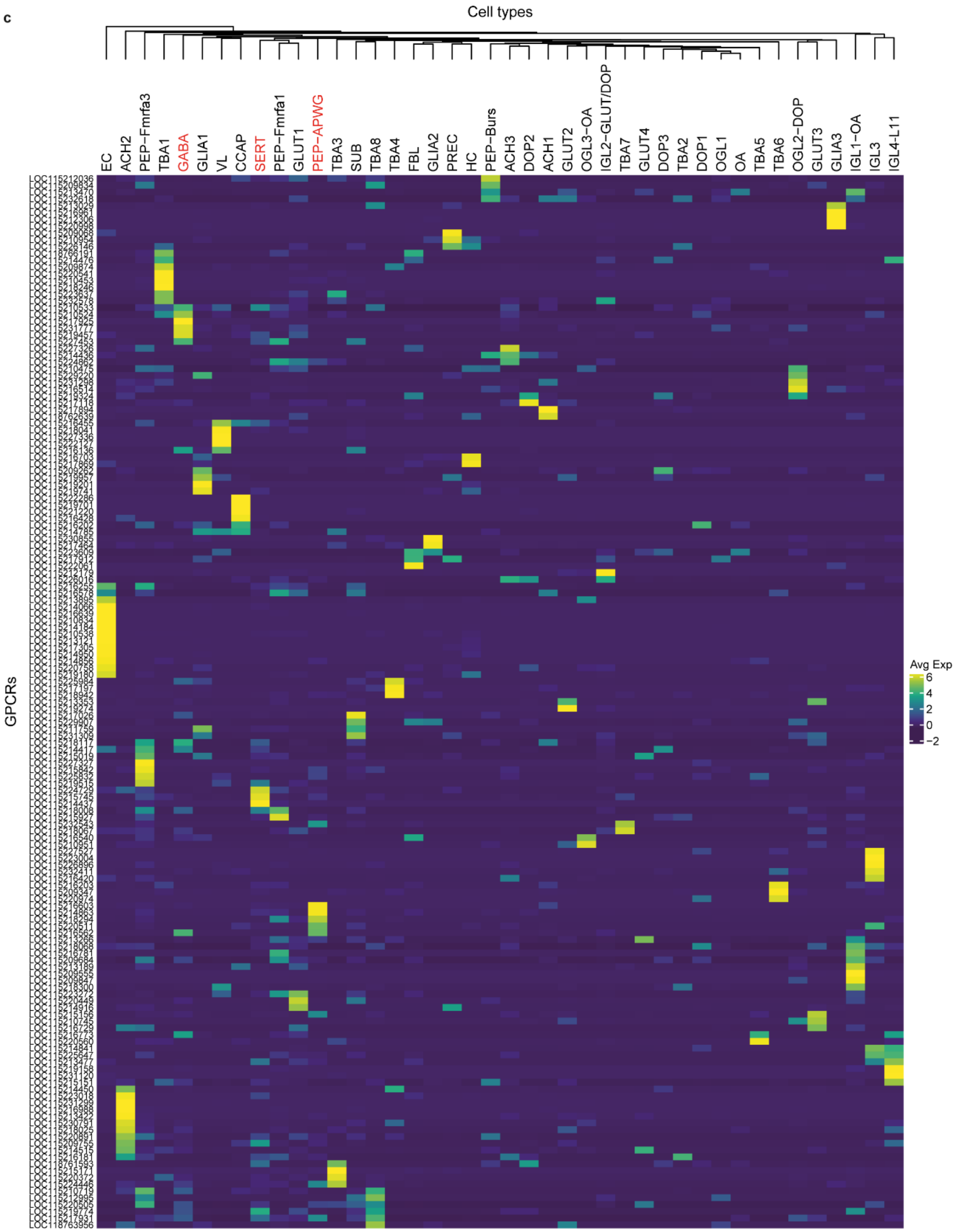


**Fig. S9 Average expression of octopus specific genes per cell type**. Scaled heatmap of differentially expressed protocadherins in **a**, Zinc-finger transcription factors in **b**, and GPCR-coupled receptors in **c**, averaged per cell type (pseudo-bulk). Significant enrichments are highlighted in red.

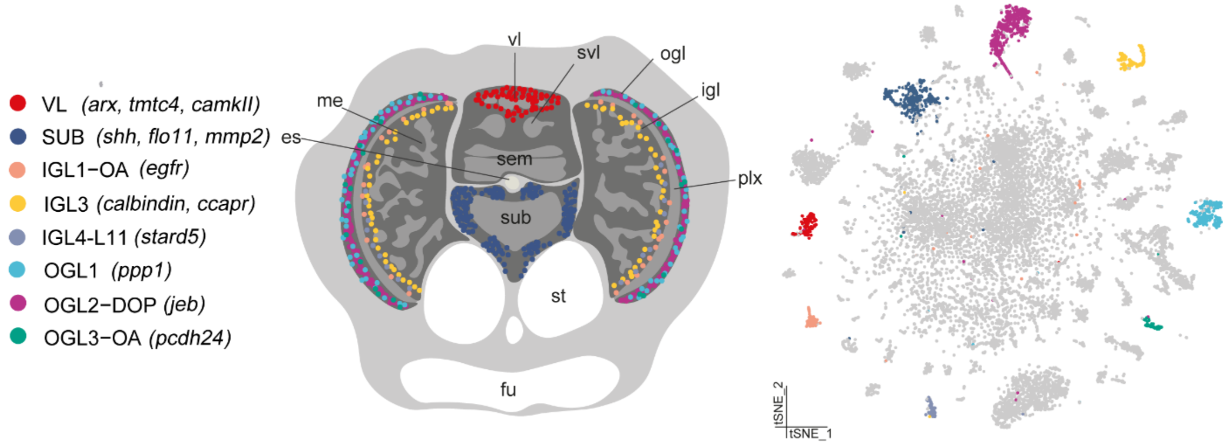


**Fig. S10 Schematic overview of distinct cell types that are spatially distributed throughout the brain.** Scheme illustrates the spatial distribution of the different cell types within the brain and within the t-SNE plot. Cell types are color coded and marker genes are visualized between brackets. DOP, dopaminergic; es, esophagus; fu, funnel; igl, inner granular layer; me, medulla; OA, octopaminergic; ogl, outer granular layer; plx, plexiform layer; sem, supra-esophageal mass; st, statocysts; sub, sub-esophageal mass; svl, subvertical lobe; vl, vertical lobe.
